## Supplementary Figures and Tables for "Ubiquitous purine sensor modulates diverse signal transduction pathways in bacteria"

to

by

Elizabet Monteagudo-Cascales, Vadim M. Gumerov, Matilde Fernández, Miguel A. Matilla, José A. Gavira, Igor B. Zhulin and Tino Krell

**Figure S1. Schematic representation of the the molecular detail of uric acid recognition by McpH-LBD.** Hydrogen bonds are shown as dotted lines and distances are indicated in Å. Hydrophobic interactions are shown as spiked arks. The amino acids that are part of the purine binding motif are labelled in red. The Figure was generated using Ligplot (1).

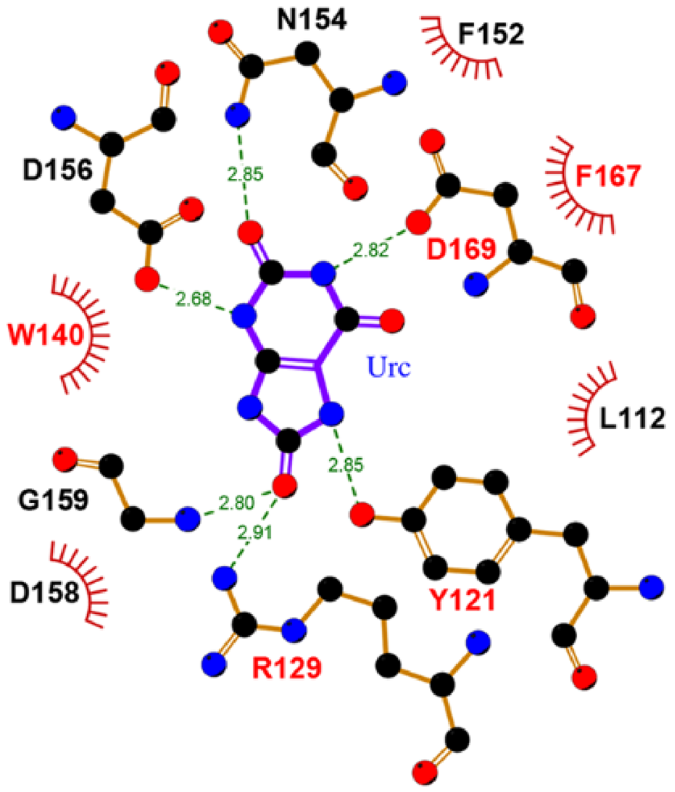

**Figure S2. Phyletic distribution of purine binding dCache_1 domains**. Blue dots indicate domains found in the RefSeq database; orange dots – domains found when the search was extended to include protein sequences from the NCBI non-redundant database. The updated taxonomy is used (2).

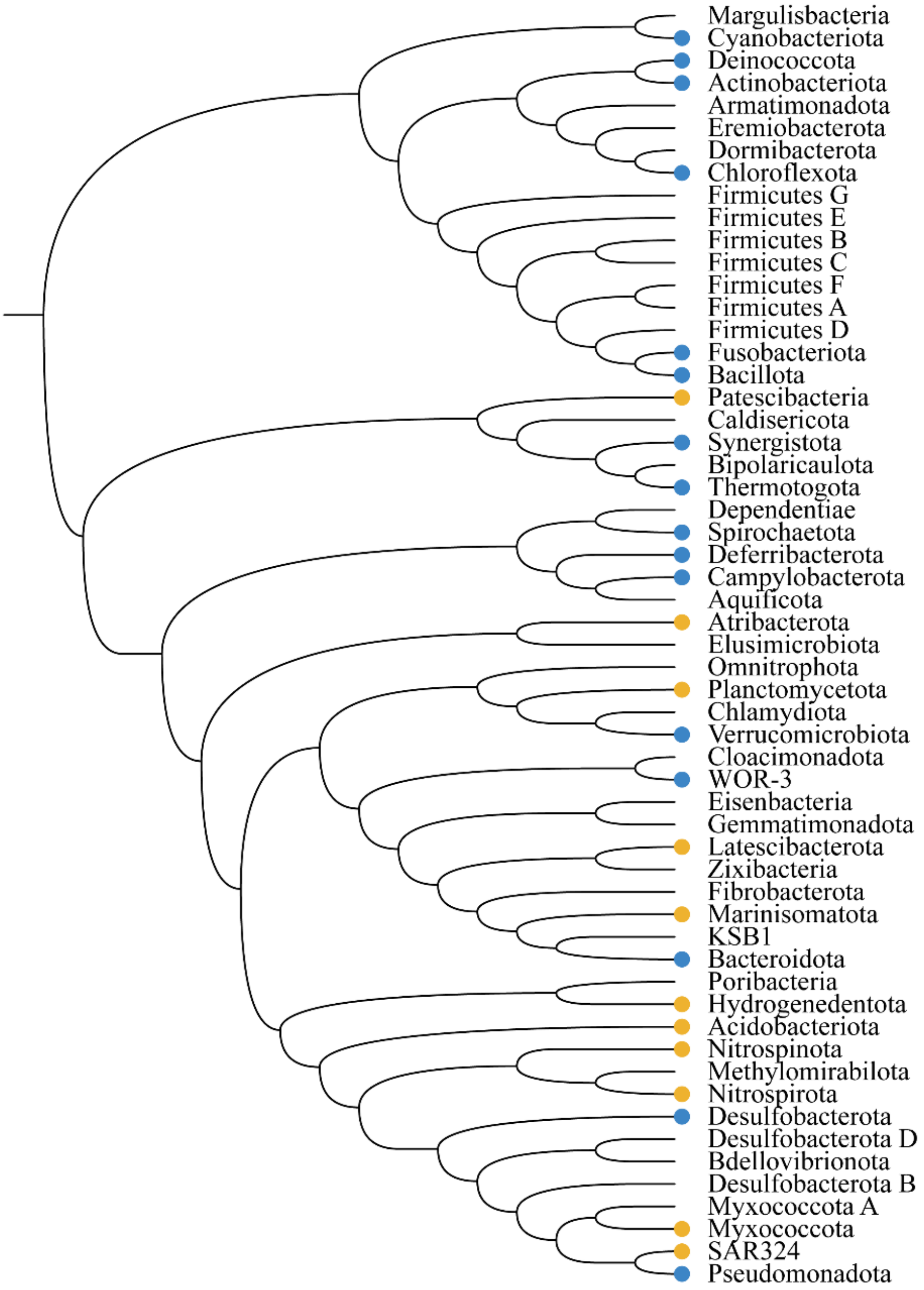

**Figure S3. Microcalorimetric titrations of the McpH-LBD mutants that were used to derive the dissociation constants as provided in Table 1.** Proteins at 9 to 18 µM were placed into the sample cell and titrated with 12.8 µl aliquots of 2 mM adenine made up in dialysis buffer. Upper panels: raw titration data. Lower panels: integrated, concentration-normalized and dilution heat-corrected peak areas and best fit using the "one–binding site model" of the MicroCal version of ORIGIN.

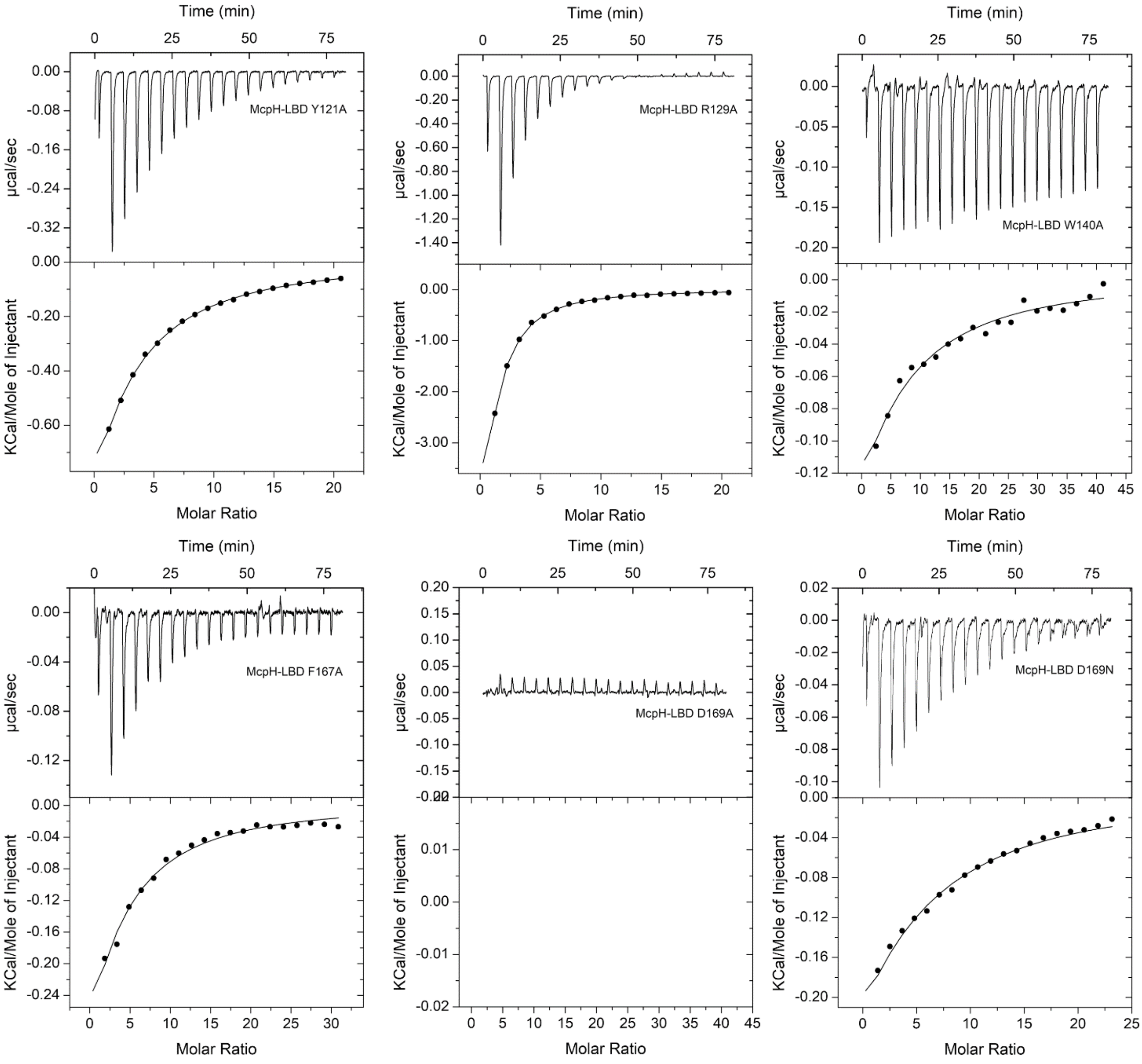

**Figure S4. Thermal shift assays of R10 and R13.** Changes in the midpoint of protein unfolding (Tm) in the presence of 2 mM ligand as compared to the buffer control. The following compounds were used (from left to right): adenine, adenosine, guanine, guanosine, inosine, xanthine, uric acid, allantoin, hypoxanthine, thymine, cytosine, purine, caffeine, theobromine and theophylline.

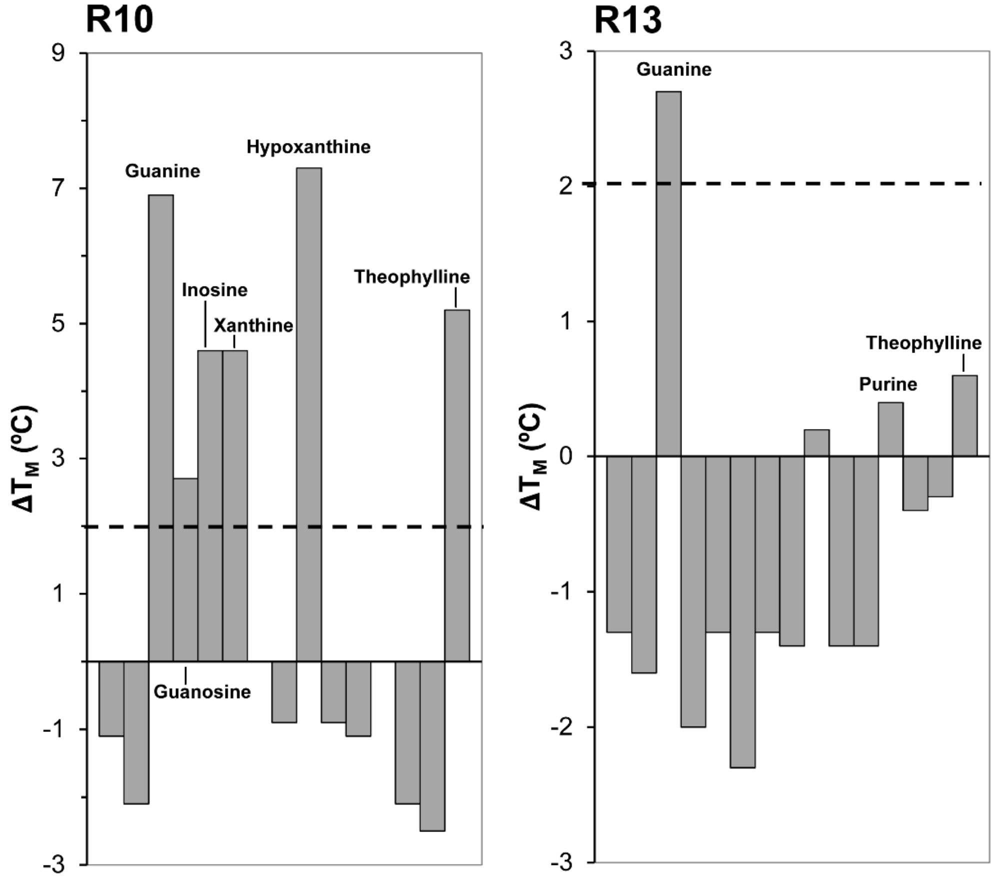

**Figure S5. Structure of ligands analysed in this study.**

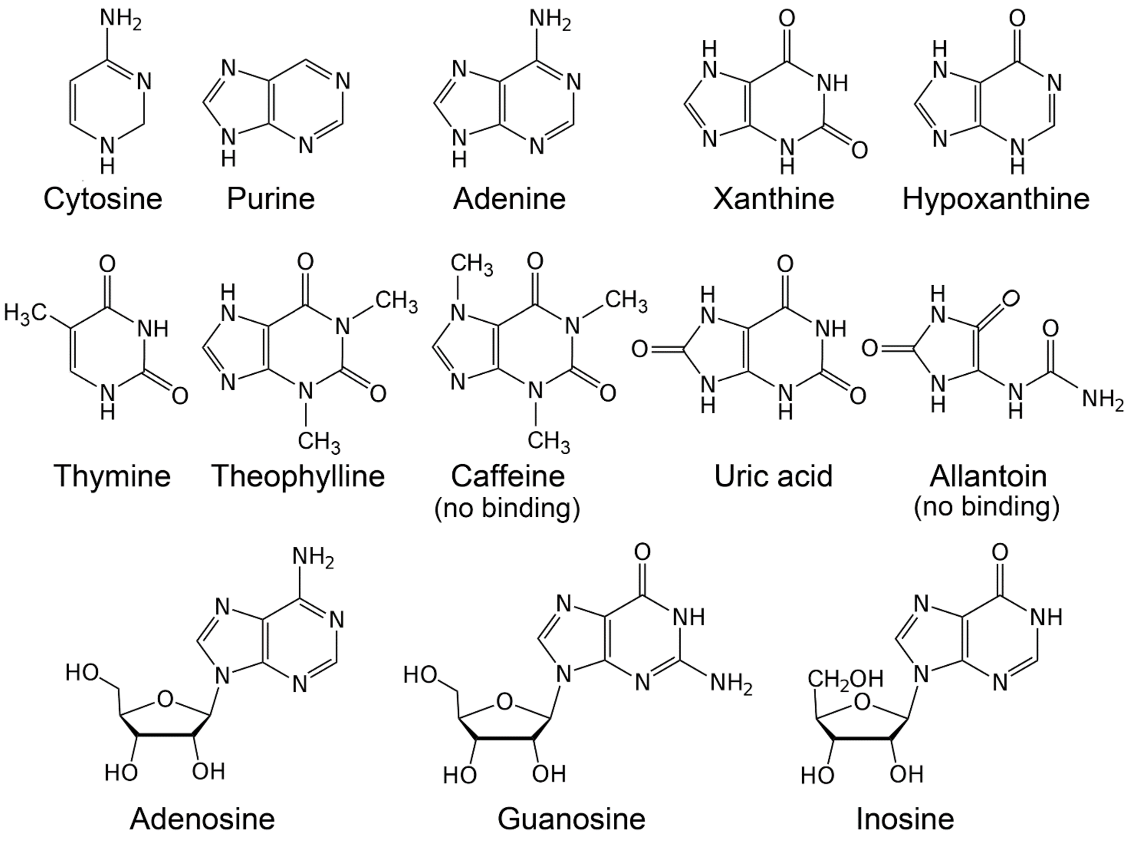

**Figure S6. Bayesian phylogenetic tree depicted in Figure 6 with proteins annotated**.

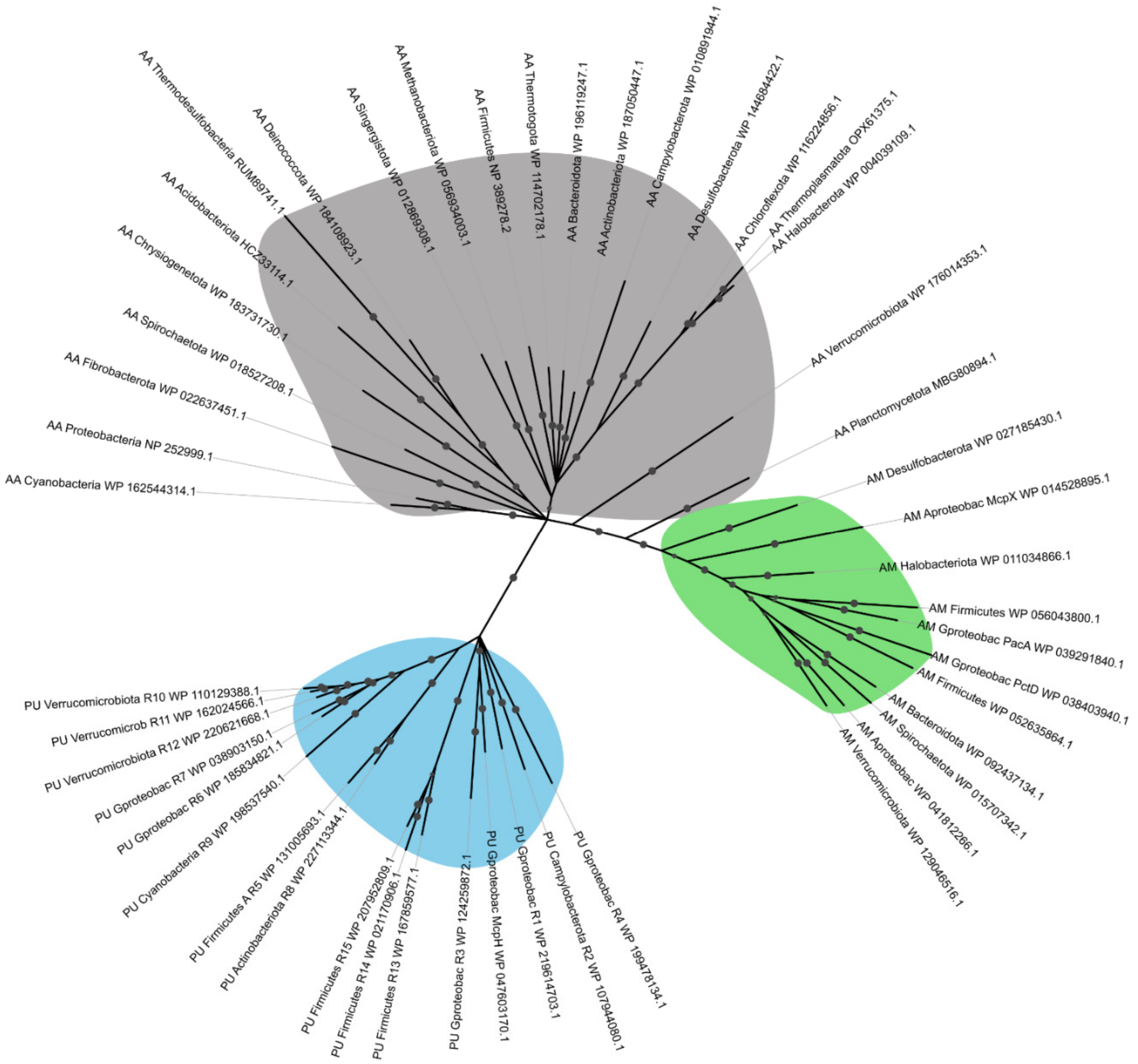

**Figure S7. Multiple sequence alignment of amino acid, amine, and purine binding dCache_1 domains.** Ligand-binding motifs are shown: purine motif (cyan), amine motif (green), amino acid motif (grey). Gproteobac: Gammaproteobacteria. Extensions R1, R2 etc. mark the proteins that have been analysed in this study (Table 2).

```
AA_Proteobacteria_NP_252999.1           ---------------YL-------QRNAIREDLESYLREMGDVTSSN-IQNWLGGRLLLV
AA_Cyanobacteria_WP_162544314.1         --------------------NDIVMREQTEERVNSNLSMAGSSVAKS-VSNWLTGRTMLT
AA_Actinobacteriota_WP_187050447.1      --------------------SYKNSIELTTQEITGSTEEQVKSMNDS-FETFLQKTENQL
AA_Bacteroidota_WP_196119247.1          ----------------------NDSKKIVVNQIKLNNYETLQNVNDYFLKNFMYDMEYII
AA_Thermotogota_WP_114702178.1          ----G------------------------KELGTTSLLSIATSGTKN-LENFLQGYVNLV
AA_Methanobacteriota_WP_056934003.1     -------------LNKMDEAMSKGMEKPVIDNGKLLATNAANLAARM-FDDYFSSIADYG
AA_Firmicutes_NP_389278.2               -------------------------KPMITEDGKNTTQNVTQSLEQN-IELQLKSYAISL
AA_Desulfobacterota_WP_144684422.1      -------------------------RKNSVENFHTGTAKELTHIEAA-MDIFMENASNIT
AA_Chloroflexota_WP_116224856.1         ------------------------YKSKQLETVQTSIDSQLALLDFS-LTNFITEVKNNV
AA_Halobacterota_WP_004039109.1         ------------------------LHEQNLDAISDNTELQLRHIEFA-LTNFISSAHYDV
AA_Thermoplasmatota_OPX61375.1          ------------------------LEDQELDTVENESITQLEHIDFA-LTNMIEESKADI
AA_Spirochaetota_WP_018527208.1         ------------------------SRTLSNRLIFQGLRQDLASLQQS-VESWAQKKLVLI
AA_Singergistota_WP_012869308.1         -------------------------HSILNDQVNTLGMEMIANNANQ-VDQYFSQLKTLT
AA_Verrucomicrobiota_WP_176014353.1     RSLVGLNAKALAQCQAIGDNAVRDSRTALIGMARTHLRDFATDQAAI-TDVKFKHIAAEV
AA_Planctomycetota_MBG80894.1           --------------------QFDTLRDTAMANARQRAQLVSESWASH-FNGQLHGIAQAV
AA_Campylobacterota_WP_010891944.1      -------------------------KTSLYESTLKNQTDLLKVTQST-VEDFRSTNQSFT
AA_Chrysiogenetota_WP_183731730.1       ----------------M-------KNHMLQQQLKNTEDLLTTVASSS-ITPWVNTRLRYI
AA_Acidobacteriota_HCZ33114.1           ---------------------------SQQQALERQAGARAQAALRH-LDEALRSDLEAS
AA_Deinococcota_WP_184108923.1          ------------------WNDRQNAERVVQAEAETTLGRLAQVAAEN-IRASLQSPTQLV
AA_Thermodesulfobacteria_RUM89741.1     -------------------LNVYQYFKIKPHFASEIMARLNRTEVNN-IKGFFAQVSEKL
AA_Fibrobacterota_WP_022637451.1        ------------------HTQHMDYDSLLTQQGKNTIHGITEHTALR-ISELLKEPALV-
AM_Desulfobacterota_WP_027185430.1      ---------------------HTSSKTTLMQEARADAANLTLASIRK-IEGTLASVEAIP
AM_Aproteobac_McpX_WP_014528895.1       --------------------LISQTQDRVETLVFDGAKTEARAIASD-IAGSVGELAAAA
AM_Gproteobac_PacA_WP_039291840.1       --------------------------------AERYLQQIAQSEALR-IQQELNYARDVA
AM_Halobacteriota_WP_011034866.1        -------------------------TTQEEKLAYQQSVEMASNYANQ-FDADMKANLAIA
AM_Spirochaetota_WP_015707342.1         ---------------------SNRSIEMAQKDAFSLAQETADKYKNA-IIAELQGARITA
AM_Verrucomicrobiota_WP_129046516.1     -----------------------RVVHSARQEANALSRTKAQAIGAE-MAHRLGRAIGTA
AM_Aproteobac_WP_041812266.1            ---------------------TTKSGSDIETLAFQSGEQLGHRYGEM-VHARLGNAMEAG
AM_Bacteroidota_WP_092437134.1          ---------------------TYNSLKTATVSSTEISNQMAVTYANQ-VVDKMDDAMSAA
AM_Firmicutes_WP_056043800.1            --------------------QLMKLYDVSLRQGELVAQNQSNAYTTK-MSIETNDALIRL
AM_Firmicutes_WP_052635864.1            --------------------SYINARNEALNAAQKRAQIVAKNYSKE-ISDELGQAITVA
AM_Gproteobac_PctD_WP_038403940.1       ------------------------------KVINERLVALARAQVSQ-IQRELEYPLTVV
PU_Gproteobac_R3_WP_124259872.1         ----------------------------KERAREQDLPTALGEIRSE-VLRQIAAPVALT
PU_Gproteobac_McpH_WP_047603170.1       ------------------------NRLTDRYLVDTALPASIEAIRND-IERMLGQPLVAA
PU_Gproteobac_R1_WP_219614703.1         --------------------QRSAQQLIETRMFEQELPNLTQRIGKE-IEKDLTSVANAA
PU_Firmicutes_A_R5_WP_131005693.1       --------------------GYQSNTNIFKNDIEHVSTLAAEGIYYQ-IDKLLSEPINVS
PU_Actinobacteriota_R8_WP_227113344.1   --------------------GYQSSRAAFEKDAERTSLLAAEGAARE-IDNRFAEPIDVS
PU_Cyanobacteria_R9_WP_198537540.1      ---------------------ANALAAARRQVVESTLPLTLDALSAD-LQQDFVQPILFA
PU_Gproteobac_R7_WP_038903150.1         ------------------------RHSLFDEISESSLPLTSDNVYSE-IQRDLLNPIFIS
PU_Gproteobac_R6_WP_185834821.1         ------------------------HDTLEEQINKDSLPLTSDNIYSE-IQQDLIRPIFIS
PU_Verrucomicrobiota_R10_WP_110129388.1 -------------------V---SRNNVRKTLAESTLPLTSDNVYSE-IQRDLLRPVFIA
PU_Verrucomicrob_R11_WP_162024566.1     ------------------------RSNMRLSITESTLPLTSDNVYSE-IQRDLLRPIFIS
PU_Verrucomicrobiota_R12_WP_220621668.1 -----------------------------------SLPLTADNIYSV-IQRDLLRPIFIS
PU_Campylobacterota_R2_WP_107944080.1   -------------------------NLYTEKVVKDELPLAVSNVAGE-IGYAIDKIINTS
PU_Firmicutes_R13_WP_167859577.1        -----------------------TQREAVDKLKTKDLLHLADSIAAK-VDGQIRKAKETS
PU_Firmicutes_R15_WP_207952809.1        ------------------------EKEVVNKLKSKDLVRIAESIASK-IDGRLQRAQESS
PU_Firmicutes_R14_WP_021170906.1        -----------------------THNAMVDKLKNRDMLYIVQSMSEK-IDGRIERAQETS
PU_Gproteobac_R4_WP_199478134.1         ------------------------MESDLKQLKEELLPNRLHSLSSR-ISEQISPLINAS

AA_Proteobacteria_NP_252999.1           E---QTAQTLAR-----------------------------------------DHSPETV
AA_Cyanobacteria_WP_162544314.1         E---FAMQSLGA-----------------------------------------KPASEMT
AA_Actinobacteriota_WP_187050447.1      D---RISKYPIMSTY--------------------------------------DKNPESI
AA_Bacteroidota_WP_196119247.1          N---DWASKDDLKNY--------------------------------RNHANQLKMVTAI
AA_Thermotogota_WP_114702178.1          D---FLSSDANVV----------------------------------GAKENKYDEVTWM
AA_Methanobacteriota_WP_056934003.1     H----MADFAVQEAY--------------------------------KNGLKGDELRQYL
AA_Firmicutes_NP_389278.2               S---RLANGELTHTFVTKP---------------------------------SKEASRLF
AA_Desulfobacterota_WP_144684422.1      R---MLAMHPEVKKIDEN-----------------LSSHTRTTRPTRPSDMTPGETEEKI
AA_Chloroflexota_WP_116224856.1         L---ALSENEIIRTQEDHD----------------FTNFLEADEE--TFEYHIGDVEQNI
```

```
AA_Halobacterota_WP_004039109.1         H---ELSLNEIVRNPDDAD----------------FTNFLNASEE--TYQYSITDEEQAI
AA_Thermoplasmatota_OPX61375.1          L---QLSMDERVQCRDDSN----------------FTNFLNATAE--DFEYNITPEEQAI
AA_Spirochaetota_WP_018527208.1         D---SVGHTLET---------------------------------------------PGM
AA_Singergistota_WP_012869308.1         L---GIASGTAELLET-------------------------------GQAVTDDDLESVM
AA_Verrucomicrobiota_WP_176014353.1     ESAAALSRELMARPHAGGNVLLYSAVNTPPLDKRQYSSLYFAPEVATETVQAAMPPLSQL
AA_Planctomycetota_MBG80894.1           D---AMAA---------------------------------------AVGQESVRNEDSL
AA_Campylobacterota_WP_010891944.1      R---ALEKDIANLPYQ------------------------------------SLITEENI
AA_Chrysiogenetota_WP_183731730.1       D---TLSYEDDL-----------------------------------LTLALERQATDTL
AA_Acidobacteriota_HCZ33114.1           Q---SLGQVVQSWW---------------------------------LEGVLEPTRPERA
AA_Deinococcota_WP_184108923.1          D-------------ITTELI---------------------------RTGQLDTADATRL
AA_Thermodesulfobacteria_RUM89741.1     LTVHDLGKNGVL----------------------------------------KFEDIINL
AA_Fibrobacterota_WP_022637451.1        ----NYTNKTYIR----------------------------------MQSLYDENDLSPL
AM_Desulfobacterota_WP_027185430.1      G---LLAFSYGK----------------------------------------NKPTASAI
AM_Aproteobac_McpX_WP_014528895.1       R---TMSGVLGR------------------------------------GHAGQSTDRAGA
AM_Gproteobac_PacA_WP_039291840.1       H---NLGQGLAA------------------------------------LPSAGIKDRAVV
AM_Halobacteriota_WP_011034866.1        R---TISTTMES---------------------------------------YETADRDEA
AM_Spirochaetota_WP_015707342.1         E---TFSTVFET------------------------------------LKDYNLTDRDMM
AM_Verrucomicrobiota_WP_129046516.1     R---TLSEALEG------------------------------------ILAEGHPSRAQA
AM_Aproteobac_WP_041812266.1            R---FIATSLVG------------------------------------LKAAGRTDREQL
AM_Bacteroidota_WP_092437134.1          R---SLAHALSG-VIG------------------------------------KNVSRQAI
AM_Firmicutes_WP_056043800.1            E---GLQQSLQQ------------------------------------MKEYNMTDRSEA
AM_Firmicutes_WP_052635864.1            E---NLGSMLKGR----------------------------------IRNENATLTRDEV
AM_Gproteobac_PctD_WP_038403940.1       H---GLANSTRLLGEPGA-----------------------------DGMPQLNASRDEI
PU_Gproteobac_R3_WP_124259872.1         R---SLATNEYILNWE-------------------------------EQGL-PEQGAAAW
PU_Gproteobac_McpH_WP_047603170.1       A---DIAGNTLLRDWL-------------------------------AAGE-DPAQAPQF
PU_Gproteobac_R1_WP_219614703.1         R---QLANDRFVLDWV-------------------------------ARGM-PKEQESIL
PU_Firmicutes_A_R5_WP_131005693.1       L---TMANDSLLKNFLDGE----------------------------KEHLNDEEFIYKL
PU_Actinobacteriota_R8_WP_227113344.1   L---AMAHDTLLADLLSAE----------------------------PTRGDDEAYADAV
PU_Cyanobacteria_R9_WP_198537540.1      S---AMAANTLLIDWV-------------------------------EQG---EQPEAAV
PU_Gproteobac_R7_WP_038903150.1         S---LMAHDTFVKDWV-------------------------------LSN---ETDPQAM
PU_Gproteobac_R6_WP_185834821.1         S---LMAQDTFVREWT-------------------------------LAG---EQDPERI
PU_Verrucomicrobiota_R10_WP_110129388.1 S---LMANDTFLRDWA-------------------------------IAG---EKDRDAI
PU_Verrucomicrob_R11_WP_162024566.1     S---LMANDTFLRDWA-------------------------------LNG---EVDINQI
PU_Verrucomicrobiota_R12_WP_220621668.1 S---MMANDAFLRDWT-------------------------------ING---EVDVKQM
PU_Campylobacterota_R2_WP_107944080.1   Y---QMTKNDYLLKWI-------------------------------DEGE-PKDGLATL
PU_Firmicutes_R13_WP_167859577.1        L---LMAEDPNVINWV-------------------------------AGGERDEALGAVV
PU_Firmicutes_R15_WP_207952809.1        L---TMAMDPELIEWL-------------------------------ASGEKDQIAGAHV
PU_Firmicutes_R14_WP_021170906.1        L---LLADDPTVLAWV-------------------------------ESGGRDEAAGEIV
PU_Gproteobac_R4_WP_199478134.1         K---LMTNDRFIADWV-------------------------------KKGA-DESRLPLV

AA_Proteobacteria_NP_252999.1           SALLEQ--------------PALTSTFSFTYLGQ--------------------------
AA_Cyanobacteria_WP_162544314.1         PEQFNV--------------PLLTSTFLMTYYGN--------------------------
AA_Actinobacteriota_WP_187050447.1      MDEFSHT-------------QSSSSEINGLYLGTE-------------------------
AA_Bacteroidota_WP_196119247.1          PKHFKPISD------QWTGYVNSIPDIAWIYLGVE-------------------------
AA_Thermotogota_WP_114702178.1          LKTFKNI-------------KDKYKDALWIYLGT--------------------------
AA_Methanobacteriota_WP_056934003.1     LNRFKQI-------------KDLNENVAFVYFGD--------------------------
AA_Firmicutes_NP_389278.2               HDDIKQI-------------KDNDDYVAMAYIGT--------------------------
AA_Desulfobacterota_WP_144684422.1      ITFLKLI-------------EASSPHLVEVFLGS--------------------------
AA_Chloroflexota_WP_116224856.1         IEILNTY-------------RTTHPYVNSVYMGR--------------------------
AA_Halobacterota_WP_004039109.1         IDILRAY-------------QISHPYVSSVYMGR--------------------------
AA_Thermoplasmatota_OPX61375.1          IDVLKDF-------------QDSHEYVNSCYMGR--------------------------
AA_Spirochaetota_WP_018527208.1         YETLRQFGPGHVNHHPFLQSSGYDDAVGTLYLAF--------------------------
AA_Singergistota_WP_012869308.1         ARYFKA--------------SAKELNVLEMYVGLE-------------------------
AA_Verrucomicrobiota_WP_176014353.1     HEMFKFM-------------FANLDSVVLAYLGT--------------------------
AA_Planctomycetota_MBG80894.1           YNFNEQL-------------IDQNDLIAGSSIAFEPG----------AWE----------
AA_Campylobacterota_WP_010891944.1      INNVGPI----------LKYYRHSINALNVYLGL--------------------------
AA_Chrysiogenetota_WP_183731730.1       MERWQTL-------------LDYSNDIFFIYFGD--------------------------
AA_Acidobacteriota_HCZ33114.1           AQLVTPL-------------LAAQRSITSLNLARADGHSLLFLRFGGAWSLRELEAP---
AA_Deinococcota_WP_184108923.1          SVTFQTM-------------LNAVPQLNGVLIGHDDG-----------------RFTSTR
AA_Thermodesulfobacteria_RUM89741.1     NRKFFPL-------------LKNQKQVTGLILADSTGREYFL-----YLDGDSWVTRVSS
AA_Fibrobacterota_WP_022637451.1        EPWFQEIARQA---------REHTPQISTVNYGNIYKNYLGI----RINDDESLNLMVQD
AM_Desulfobacterota_WP_027185430.1      STDLLGF-------------ILFNSAVYGSCVAYEPY----------AFD-RDVEF----
AM_Aproteobac_McpX_WP_014528895.1       INLLKAN-------------LEQHGFAFGSWFAEEPK----------AYDGKDVIDNTER
AM_Gproteobac_PacA_WP_039291840.1       DKMMEYA-------------LRDNPEYLSISVIFEEN----------VFDGRDAEFADQP
AM_Halobacteriota_WP_011034866.1        LLILENL-------------LRDNPHLLGTYVAFEPD----------AFDGKDAEYTNSP
AM_Spirochaetota_WP_015707342.1         NDILKNA-------------LANKEYITAFCIAYDPN----------AMDGKDAQYAGQG
AM_Verrucomicrobiota_WP_129046516.1     DAMLRGS-------------LEGNTDYIGVWTLWEPN----------AFDGRDADYVNKP
AM_Aproteobac_WP_041812266.1            SIWLKSI-------------AEANPDFLGVWVGMEPN----------ALDGRDAEFANKP
AM_Bacteroidota_WP_092437134.1          QQMAGSI-------------LLGDEDFLGYTVCFEPN----------AYDAKDAFFANKP
AM_Firmicutes_WP_056043800.1            VRLIENF-------------VREQPYILGVFTVWEPN----------AFDNQDGNFRNKS
AM_Firmicutes_WP_052635864.1            SEIFKNA-------------LADNPQFIGISIAFEPN----------AFDSLDAQFDGDS
```

```
AM_Gproteobac_PctD_WP_038403940.1       SALLRST-------------VQNNPKLLDTFMAWEPN----------AFD-TDAAFAGQP
PU_Gproteobac_R3_WP_124259872.1         KTYAQTL-------------KNESHAATIAWVSE--------------------------
PU_Gproteobac_McpH_WP_047603170.1       IEYLTAA-------------KQRNHAFTTLFAST--------------------------
PU_Gproteobac_R1_WP_219614703.1         IDQLKDM-------------TAQYGLVTASFADR--------------------------
PU_Firmicutes_A_R5_WP_131005693.1       QDYLNVY-------------RNKYSYDSVFLVST--------------------------
PU_Actinobacteriota_R8_WP_227113344.1   CAYLESY-------------REAFGFDSVFLVST--------------------------
PU_Cyanobacteria_R9_WP_198537540.1      QRYLSRV-------------QAQHGATTVFFVSE--------------------------
PU_Gproteobac_R7_WP_038903150.1         TRYLREI-------------DRRFNTVVSFFVSN--------------------------
PU_Gproteobac_R6_WP_185834821.1         IRYLREI-------------QRQYQTISSFYISD--------------------------
PU_Verrucomicrobiota_R10_WP_110129388.1 VRYLHEI-------------KIKYGTVSSFFVSD--------------------------
PU_Verrucomicrob_R11_WP_162024566.1     TKFLHEI-------------KVEYSTISSFFVSD--------------------------
PU_Verrucomicrobiota_R12_WP_220621668.1 QRYLEEI-------------RREYGTVTSFFISE--------------------------
PU_Campylobacterota_R2_WP_107944080.1   FNYNTDL-------------MKAFNLSTAMFVSD--------------------------
PU_Firmicutes_R13_WP_167859577.1        GRKLTLL-------------TSEHGYDNSFLVSA--------------------------
PU_Firmicutes_R15_WP_207952809.1        LQKLDNL-------------AKGFDYSNGFIASR--------------------------
PU_Firmicutes_R14_WP_021170906.1        KTKITDI-------------GKNYDYVKAFVAST--------------------------
PU_Gproteobac_R4_WP_199478134.1         AEELNSI-------------KQLSGSDSTFYVVNM-------------------------

AA_Proteobacteria_NP_252999.1           ----------------------------QDG-VFTMRP-----------DSPMPAGYDP-
AA_Cyanobacteria_WP_162544314.1         ----------------------------SEG-QFFQWP-----------PSTLPEGYDP-
AA_Actinobacteriota_WP_187050447.1      ----------------------------KDG-KTLIFP-----------KADLPADFDP-
AA_Bacteroidota_WP_196119247.1          ----------------------------EDG-SIFITP----------IDPTMPDDYDC-
AA_Thermotogota_WP_114702178.1          ----------------------------SDK-KFYIYP-----------DAELPEGYDP-
AA_Methanobacteriota_WP_056934003.1     ----------------------------ENG-NMYLWP-----------DEPLPPDYDP-
AA_Firmicutes_NP_389278.2               ----------------------------AKK-EMFTYP-----------KADFAEDYDP-
AA_Desulfobacterota_WP_144684422.1      ----------------------------RWG-GFTYAG-----------EDALPAGYDP-
AA_Chloroflexota_WP_116224856.1         ----------------------------ENG-SFVRSH-----------PRNEPTQYDP-
AA_Halobacterota_WP_004039109.1         ----------------------------ENG-AFVRSY-----------ERARPTAYDP-
AA_Thermoplasmatota_OPX61375.1          ----------------------------ENG-AFVRAY-----------PRASPTQYDP-
AA_Spirochaetota_WP_018527208.1         ----------------------------RDG-VHTTGS-----------AWNIPDDYDP-
AA_Singergistota_WP_012869308.1         ----------------------------STG-KVGTGG-----------DWVEKPDYDA-
AA_Verrucomicrobiota_WP_176014353.1     ----------------------------VDG-VELQYP-----------WSHTPLAYDP-
AA_Planctomycetota_MBG80894.1           ----QETDRYAPYSWI------------TDG----------S-----TNRRDLALDYDY-
AA_Campylobacterota_WP_010891944.1      ----------------------------NNG-KVLLSQKSND-----AKMPELRDDLDI-
AA_Chrysiogenetota_WP_183731730.1       ----------------------------REG-NVTMLP----------DDEPLPPDFNM-
AA_Acidobacteriota_HCZ33114.1           --GPGARIRW--------------RRLDQNG-QVLSNE------------PWTPMAYDP-
AA_Deinococcota_WP_184108923.1          RGGSSDAGRSLSTIETRPTRRATVTVLDERG-RIVSRR-------------VDTGNYDP-
AA_Thermodesulfobacteria_RUM89741.1     V---------------------------KGSVTHMVFQKWNGPDEPVKKWEKNVRDYDP-
AA_Fibrobacterota_WP_022637451.1        TH--------------------------TND-SLYIYETLHRS----GPIAAVFGAYDP-
AM_Desulfobacterota_WP_027185430.1      ---------FAPYAYM------------PGG-RPMFTY--------------LSADYNY-
AM_Aproteobac_McpX_WP_014528895.1       --GGNADGAFTPYWSKD-----------RNG-NIQLST------------------FKAD
AM_Gproteobac_PacA_WP_039291840.1       --GQAPKGRYAWFVDRD-----------QAG-NYAMHP---------------LLSYLTP
AM_Halobacteriota_WP_011034866.1        --AHDGTGRFVPYWNK------------MNG-TASVAP---------------LLHYDS-
AM_Spirochaetota_WP_015707342.1         P-AYDETGRYAPYWNK------------LGG-NIDVEF---------------LPDIDS-
AM_Verrucomicrobiota_WP_129046516.1     --GHDATGRYIAYWNR------------GSG-KVIVEP---------------LVDYTTE
AM_Aproteobac_WP_041812266.1            --GSDASGRFLPYWNR------------GSG-TVALES---------------LVGYDEP
AM_Bacteroidota_WP_092437134.1          --GHDNTGRFVSYMTKN-----------GSG-GFVVEP---------------LVDYENE
AM_Firmicutes_WP_056043800.1            S-YDDDTGRFVPYIVR------------QGD-KIVAYP---------------NKNYENI
AM_Firmicutes_WP_052635864.1            --RYYEKGQFATYFVRGN---------LNSG-KDMYST------------ITQEPLRDL-
AM_Gproteobac_PctD_WP_038403940.1       GKGYGPDGRYLPWWYRG-----------ADG-KPIVEA--------------MADSIDSE
PU_Gproteobac_R3_WP_124259872.1         ----------------------------KTG-KYLDEN-----------GFSRTVQRSD-
PU_Gproteobac_McpH_WP_047603170.1       ----------------------------ETG-HYYNEN-----------GLDRTLSRSN-
PU_Gproteobac_R1_WP_219614703.1         ----------------------------QSA-AYYNQD-----------GFLRNLTP---
PU_Firmicutes_A_R5_WP_131005693.1       ----------------------------KTN-NYYHFN-----------GLDRNLSANN-
PU_Actinobacteriota_R8_WP_227113344.1   ----------------------------ESN-RYYHFN-----------GVDRTLERDN-
PU_Cyanobacteria_R9_WP_198537540.1      ----------------------------ASR-RYYHPT-----------GILKTVSPGS-
PU_Gproteobac_R7_WP_038903150.1         ----------------------------NTH-RYYDPE-----------KISHTLLETS-
PU_Gproteobac_R6_WP_185834821.1         ----------------------------RTG-HYYHYT-----------GILKQVSESN-
PU_Verrucomicrobiota_R10_WP_110129388.1 ----------------------------KTL-KYYYAH-----------GLLKTVSEDE-
PU_Verrucomicrob_R11_WP_162024566.1     ----------------------------ISH-NYYHAQ-----------GLLKEVKENE-
PU_Verrucomicrobiota_R12_WP_220621668.1 ----------------------------KSR-NYYYWG-----------GVLKQVDENE-
PU_Campylobacterota_R2_WP_107944080.1   ----------------------------KTL-NYYTND-----------KILKQLSKDN-
PU_Firmicutes_R13_WP_167859577.1        ----------------------------VSN-RYWGEG----------GTVLRTMSRSD-
PU_Firmicutes_R15_WP_207952809.1        ----------------------------MTG-SYWIEG----------KQMSKILSQTN-
PU_Firmicutes_R14_WP_021170906.1        ----------------------------VTN-QYWEDN-----------KVVKPLSKTS-
PU_Gproteobac_R4_WP_199478134.1         ----------------------------KSGLEFLGYD----------QKFFRTPLADY-

AA_Proteobacteria_NP_252999.1           -------RSRPWYKDAVAA---GGLTLTEPYVDAA-TQE--LIITAATPVKA-A---GNT
AA_Cyanobacteria_WP_162544314.1         -------RKRPWYQAAASF---GKLTLTEPYIAAS-SGE--LNITAAAPRYQ-G---GQL
AA_Actinobacteriota_WP_187050447.1      -------RQRDWYQSALKQ--KNKTIWTEPYTDQA-TNE--LVITAAKAIYDDR---DEL
AA_Bacteroidota_WP_196119247.1          -------RTRSWYKATVNN--NEKIIWTEPYVDAGDLGN--VIVTVAKAIHK-N---DSL
```

```
AA_Thermotogota_WP_114702178.1          -------TKRPWYVDAVKN--KGKVIITEPYLDAS-TSD--IVITVAKAVVN-N---GQI
AA_Methanobacteriota_WP_056934003.1     -------RVRPWYIKAKEN---NGPSYTEPYRDAF-TGK--WVITYSEPVYV-D---GKF
AA_Firmicutes_NP_389278.2               -------TSRPWYKLAAET--PDQVVWTEPYKDVV-TGD--MIVTASKAILD-R---QKV
AA_Desulfobacterota_WP_144684422.1      -------RSRPWYQTALAN--PARTSITSAYMST--TGE--PVISLMHGIQSER---NEL
AA_Chloroflexota_WP_116224856.1         -------RDRPWYILAKEN--PGTVMITEPYQSLT-TSD--INVGIVTALVDQT---GEV
AA_Halobacterota_WP_004039109.1         -------RERPWYILAKEH--PGEVMVTEPYSSVT-TPD--VNIGTVTALLDED---GEV
AA_Thermoplasmatota_OPX61375.1          -------RERPWYQLAMET--PGEVRITDPYSAVT-TDD--VNIGIVLALMD-N---GTV
AA_Spirochaetota_WP_018527208.1         -------RERGWYQTAQSQ---TGPGFSDSYVDAH-TGN--LVITAARPLRDQE---GEI
AA_Singergistota_WP_012869308.1         -------RKRPWYIQAVQE---DKVILTAPYVDAN-TGG--LVITVATPVKSTS---GKL
AA_Verrucomicrobiota_WP_176014353.1     -------RQRGWFKSAAAH--PGETCWSAPYASASLDNK--LVISCSRAIQDAD---GKL
AA_Planctomycetota_MBG80894.1           -------LERDWYRVGVD----GEAGWTEPYDGPV-FGS--LLVTYSSPVIQ-N---GKV
AA_Campylobacterota_WP_010891944.1      -------KTKDWYQEALKT---NDIFVTPAYLDTV-LKQ--YVITYSKAIYK-D---GKI
AA_Chrysiogenetota_WP_183731730.1       -------FERPWYLRATES--PGQIAWTDLYDEII-TGI--PQISAVVTVHDPE--SLEP
AA_Acidobacteriota_HCZ33114.1           -------RTRPWYLAGAAA---PAPTWTEPYAFYT-TQD--PGITYTLPVRD-T---TGL
AA_Deinococcota_WP_184108923.1          -------RTRPWYTLAVAT--PGITVWTPPYTFAS-SGQ--PGVTVARAVDA-G---ADG
AA_Thermodesulfobacteria_RUM89741.1     -------RKRPWFRSSSSE---RRVFWSPVYRFYS-TGK--PGVTASVSWAESG--NPPR
AA_Fibrobacterota_WP_022637451.1        -------TQRPWFAPLFES---PTKQWSEVYVNLDEKAE--LTITHMLPVFSPQ---NTL
AM_Desulfobacterota_WP_027185430.1      -------PQADWFLIPKEI---RRPIWSEPYFDEG-GGN-VVMSTYSIPFFREEDGRKRF
AM_Aproteobac_McpX_WP_014528895.1       -------YAAEWYGLAAKS---GKGAITQPYLAEG-TDVPTTMTSIAYPVMS-N---GRM
AM_Gproteobac_PacA_WP_039291840.1       -------GQGDYYLLPQKS---QKDTLIEPYTYAY-NGVPTLLTSVAAPIVS-Q---GKL
AM_Halobacteriota_WP_011034866.1        ---------SDYYQLPKAT---EKDVLTEPYFYEG-----VFMVSYVSPIMK-E---GEF
AM_Spirochaetota_WP_015707342.1         ---------EDWYIVPKAE---RHEYITDPYPYGL-QGRTVMLASLIFPIIH-S---DKF
AM_Verrucomicrobiota_WP_129046516.1     -------GAGDYYLLAKHS---NQETVLEPYIYKV-AGRDVLMTSLVVPVNRAD---GTF
AM_Aproteobac_WP_041812266.1            G------SDGAYYQIPKRT---GHAMVVEPYSYTV-AGRKVLMVSMSVPIVE-N---GRV
AM_Bacteroidota_WP_092437134.1          -------SAAPWYWIPMRT---MKEFVTEPLMYPI-QGKNVYMVSFMCPIIT-N---GKF
AM_Firmicutes_WP_056043800.1            -------GDGDYYQIPKRT---KKFALMEPYYYDI-NGERILISSFVYPILDEQ---GKF
AM_Firmicutes_WP_052635864.1            -------EISDYYIVPKQT---LSNVMIEPYIYEV-QGKEVLMTTCSSPIVI-D---GKY
AM_Gproteobac_PctD_WP_038403940.1       KLLPTGVRENEFYACPKEN---KRPCIIDPAPYEM-GGKTVMMSSFNVPIMV-G---DQF
PU_Gproteobac_R3_WP_124259872.1         -------ATDNWFYDFLSK--GKQLEIKLGKDKAS--SV--YNLFIDARFDV-N---GK-
PU_Gproteobac_McpH_WP_047603170.1       -------PKDKWFYGYIDS--GAERFINIDIDGAT--GE--LALFIDYRVEK-E---GKL
PU_Gproteobac_R1_WP_219614703.1         -------EQDAWFYGYTKS--PQDLMLSI-FRETN--GE--VKLFVNFQQLN-----GR-
PU_Firmicutes_A_R5_WP_131005693.1       -------SENQWYYTFLKN--DDEYSLNVDNDEAS-NNS--ITVFVNCKIKDDN---GAT
PU_Actinobacteriota_R8_WP_227113344.1   -------PENTWYFDFLDR--DEAYSLNVDNDEAT-ADE--ITMFVNARILDEQ---GST
PU_Cyanobacteria_R9_WP_198537540.1      -------AQDAWFFRLRAS--ASSYEVNLDRDTAD-PSR--TTVFVNYKLLGDG---GRF
PU_Gproteobac_R7_WP_038903150.1         -------PEDKWFFDIRDEKDGDPYDIEIGVDPEN-RTR--MDIFINYKVFDYS---GNF
PU_Gproteobac_R6_WP_185834821.1         -------PNDAWFYRVKNSAPDKNFEVNIDIDTAN-SQQ--TVVFVNYKVFDFE---NRF
PU_Verrucomicrobiota_R10_WP_110129388.1 -------PRDEWYFRVREM--DEPYEINVDPDMAN-QDA--LTIFINYRVRDYA---GNF
PU_Verrucomicrob_R11_WP_162024566.1     -------PRDAWYFKARDM--DAPYEINVDLDMAN-QDQ--MTIFINYRVFDYN---DNF
PU_Verrucomicrobiota_R12_WP_220621668.1 -------PRDVWYFRVREM--EKPFEINVDIDMAN-NDA--LTVFVNYRVYDYE---GNY
PU_Campylobacterota_R2_WP_107944080.1   -------PRDSWYFDVKNG--KEVNSLNIQVSEAT--GS--LTLYVNSKVEK-D---GKF
PU_Firmicutes_R13_WP_167859577.1        -------PEDAWFYANVDN--RKPIEIVLDYAIAR--GD--TFVFVNALIGG----AERA
PU_Firmicutes_R15_WP_207952809.1        -------PADKWFYDSFAS--QKRMALNIDYNDSR--RD--TFVFVNVLVGD----VESP
PU_Firmicutes_R14_WP_021170906.1        -------YTDKWFYKALKS--GKKIELNIDYDSIH--NE--TFIFFNTLVGD----VRQP
PU_Gproteobac_R4_WP_199478134.1         -------PYKEFYPNFLAK--NKDYELNLQYADQK--------LYINYRSREMDPTTGKP

AA_Proteobacteria_NP_252999.1           LGVVGGDLSLKTLVQIINSLDF--S-GMGYAFLVSGDGKILVHP-DK----------EQV
AA_Cyanobacteria_WP_162544314.1         VGVVGSDFSIQMLDQMLKDANL--G-DLGFLFITDKSGKVLIHA-NQ----------DLI
AA_Actinobacteriota_WP_187050447.1      IGVLGVDISIDTLITMVNQTKF--G-ETGYTVLLDQKGSFVTHP-DK----------EKI
AA_Bacteroidota_WP_196119247.1          VGVIGMDIKLKKFSNIINNIQF--G-ENGYLMLLSKKGDVYAHP-NN----------KML
AA_Thermotogota_WP_114702178.1          IGVVALDFKASELANSLLSNKF--G-ENGYSYLLSSDGKTLLHV-DE----------EKI
AA_Methanobacteriota_WP_056934003.1     VGVVGIDVFVSTLMKEAMDIKL--G-KSGYIVLVNQEGLVLMHP-KE----------EYI
AA_Firmicutes_NP_389278.2               IGVASYDLKLSAIQSMVNKQKV--P-YKGFAFLADASGNLLAHP-SN----------Q--
AA_Desulfobacterota_WP_144684422.1      VGIAGLDVSLGALTSLIQRSQI--G-ETGYVMLVQGDGVILANPRNP----------ETN
AA_Chloroflexota_WP_116224856.1         YGVIGADVTLTNLTEFISGFDV--G-HSGQLLLVNEQGAILANK-NS----------QLL
AA_Halobacterota_WP_004039109.1         YGVIGADITLVNLTEYLASIES--I-GNQEMILTDRSGTILASY-NT----------TVL
AA_Thermoplasmatota_OPX61375.1          YGVVGADITLVGLTDYLRQASE--V-SGRDMLLSDDRGTILAYK-QE----------QLL
AA_Spirochaetota_WP_018527208.1         FGVLATDISLGQITEAISEYCL--G-EGGYAFLVNQEGSLLAHP-DT----------EXA
AA_Singergistota_WP_012869308.1         LGVAGIDVDLKTLSDMITSYKV--F-GKGYGFLLDREGNMICHP-KS----------EMI
AA_Verrucomicrobiota_WP_176014353.1     FGVLALDISVEAVIHDFISTQE--M-RRGSAFLLNDSGEIIAQE-GM----------DLL
AA_Planctomycetota_MBG80894.1           IGVMAADIALLPLQQRLKEEAP----EGLSTILASASGRLIAHP-NP----------ALI
AA_Campylobacterota_WP_010891944.1      IGVLGVDIPSEDLQNLVAK-------TPGNTFLFDQKNKIFAAT-NK----------ELL
AA_Chrysiogenetota_WP_183731730.1       VGVMALDVSLHQLRRIVMDVEL--P-SDAELVLLDRNHQLIVSSTGK-------------
AA_Acidobacteriota_HCZ33114.1           RGVAALDFLLDDLTAQVWTAQP--T-SRSRCLVVDAQGRALVLPRDPAFETPEGRRRAFL
AA_Deinococcota_WP_184108923.1          TVVVGADVQLRQVASFLQGVQI--G-GHGRAFVTDAQGHVIATS-PT---WPG----NVT
AA_Thermodesulfobacteria_RUM89741.1     TVVFALDITLSQIQKFLELQDD--N-KHELLFILNPRDKFFITP-------------KYI
AA_Fibrobacterota_WP_022637451.1        YGAAIVDVKLGRLHAFLQEEEI--P-PGGAVYITTSEGSLLASSYKT---------QDTI
AM_Desulfobacterota_WP_027185430.1      LGVVTADISLEWLRTFIKSISI--Y-QSGYAFLLSRNGVFLSHP-ND----------QFI
AM_Aproteobac_McpX_WP_014528895.1       IGVSGVDISLAALADRLSAVKP--F-GSGRVYLLSQSGKWLAAP-IP----------ELL
AM_Gproteobac_PacA_WP_039291840.1       WGVVTSDISLASLQQKINQIKP--WEGGGYAMLLSSAGKVISYP-DK----------SQT
AM_Halobacteriota_WP_011034866.1        AGIGGVDVSLEYVDEVVSKVRT--F-DTGYAFMVSNSGVILSHPTHK----------DWI
AM_Spirochaetota_WP_015707342.1         IGIISSDIVLDKLQEMVDKVNPH-G-QEGYTEIISHSGAVIAHP-NK----------DYL
```

```
AM_Verrucomicrobiota_WP_129046516.1      AGVVGVDLPLETLGAEIAKVKV--G-ETGYAALVSNTGIYAAHP-RA----------ERL
AM_Aproteobac_WP_041812266.1             IGVAGIDLSTDGIWSMLKTVKP--F-DSGSIHLISNDGVWAGHP-DS----------ERM
AM_Bacteroidota_WP_092437134.1           VGVTGVDLSINYLQDMVVKANV--FDGHGNFDIVSHQGVFAANSGNP----------DFV
AM_Firmicutes_WP_056043800.1             LGVVGADISLDMVQQEVEKIRP----MGGYATMITAGDSYLANGFDR----------ALV
AM_Firmicutes_WP_052635864.1             YGVVGVDIEVDFIKELVKGSEND-M-EFKDILIISSKGNIVGS--KY----------EMA
AM_Gproteobac_PctD_WP_038403940.1        RGAVGADLSLAFIQDLLKRADQQLYDGAGEMALIASNGRLVAYTRDD----------SKL
PU_Gproteobac_R3_WP_124259872.1          IGVAALGLSVNELADFIRKQKI--G-NSGFVYLVSPDGAFVIHR-DA----------ALA
PU_Gproteobac_McpH_WP_047603170.1        VGVAGMGLRMTELSKLIHDFSF--G-EHGKVFLVRNDGLIQVHP-DA----------AFS
PU_Gproteobac_R1_WP_219614703.1          -GLAGLAKSLDSMVSMLANFRI--G-DSGFVFMTDGSGKVKLHP-DA----------ARI
PU_Firmicutes_A_R5_WP_131005693.1        MGIIGVGLKVNSLQMLLKGYND--K-FDVVARLIDDKGFVQLAV-DK----------TGH
PU_Actinobacteriota_R8_WP_227113344.1    LAIVGVGFRMDDLKELLAGFEA--R-TDTRVRLLADDGTIRAST-DP----------NEN
PU_Cyanobacteria_R9_WP_198537540.1       LGAVGLGRSTSQLTRRIQQAER--T-NGIQVMFLDGRGRILFSP-RR----------GQA
PU_Gproteobac_R7_WP_038903150.1          IGVTGVGLPVQRVTQLIETYEQ--R-YNRTIYLIDEDGDVMLHS-KA----------FHR
PU_Gproteobac_R6_WP_185834821.1          LGVIGVGLSSDAVSALVEKYQK--R-YNRHIYFINELGEVTLHG-SH-------------
PU_Verrucomicrobiota_R10_WP_110129388.1  IGATGVGLTVTKVNRLISRYEA--K-YDRQIYFVDASGNVVLRP-SN----------STM
PU_Verrucomicrob_R11_WP_162024566.1      IGMTGVGLTVKNVNNLISHYEA--K-YQRQIYFLNKDGEIVLRP-SN----------SPL
PU_Verrucomicrobiota_R12_WP_220621668.1  IGAAGTGLTVNRVNALIEEYEG--R-FNREIFFVDREGNIILGP-SK----------GRL
PU_Campylobacterota_R2_WP_107944080.1    YGVSAIGMNLDDIVNLVTSKTM--G-EGSKFLMVDSSGIVKIEK------------SDRV
PU_Firmicutes_R13_WP_167859577.1         LGETGVGLSLKQSAEQFQQFKY--G-DKSHLWLVDREGTIYLSD-RY----------EQA
PU_Firmicutes_R15_WP_207952809.1         VGVAGIGLSLKELANNFTNYKY--G-ANSNVWLVDKSGKIYLSD-RV----------EDI
PU_Firmicutes_R14_WP_021170906.1         VAVAGVALSLGDIAKEFGSYKF--G-EHSNLWLVDKQGKIHLAD-DL----------EYN
PU_Gproteobac_R4_WP_199478134.1          LVVAGLAIKVDKLIDMVKQLTI--G-KSGRAMLVTDQGVIQAKG-ESP-------AIDLI

AA_Proteobacteria_NP_252999.1            MK---------TLSEVYPQNT--------------PKIATG------FSEAELHGHTRIL
AA_Cyanobacteria_WP_162544314.1          GK---------TLTDIYPGAQ----------------VSTG-A----MQDVESADGARLL
AA_Actinobacteriota_WP_187050447.1       QQ---------DISKENIFKK--------------MKGESG------SMIEEFEGKSRIV
AA_Bacteroidota_WP_196119247.1           TK---------NISNKEWVKT-------------ILSNEKG------TDIHTWNNEKVVI
AA_Thermotogota_WP_114702178.1           GK---------SVADMDWFKQ-------------MINSKNKSG------VIEYVYDGIKRMA
AA_Methanobacteriota_WP_056934003.1      NK--------LNIYKVPELQE------LA--AELKKGKQEG------TVIYTFEGIRRIA
AA_Firmicutes_NP_389278.2                GK---------NISKDQTLQT-------------IASEKKG-I---------QDVNGKMV
AA_Desulfobacterota_WP_144684422.1       FK---------KLTESGIEAF-----------RVLDKTDRG------SVSLEIKGKEWLA
AA_Chloroflexota_WP_116224856.1          FK---------NVQTMLGESS-----------VEFMKEDHG----------IFTYGRNYC
AA_Halobacterota_WP_004039109.1          FE---------DIGTVLGEQA-----------GFFLATEEG----------VLVHDGRYL
AA_Thermoplasmatota_OPX61375.1           FT---------NVSAVLGDDT-----------DDFLGTEQG----------RIRLENDFF
AA_Spirochaetota_WP_018527208.1          GT---------SLEEIDGYAEV---------ARGILGKGSG-ESGGRQFSTTIAGRPHII
AA_Singergistota_WP_012869308.1          MK--------ENITKPSGLIT-PELAEAG---RKMISGSPG------FVDYSFQGELRRT
AA_Verrucomicrobiota_WP_176014353.1      GQGWN---LQVDLVNLLHSEE-TQLQQLA---AEMIALNSG------CLRLPLGDTVYYV
AA_Planctomycetota_MBG80894.1            LS--------DSLQSEADRTLTPELHAWV---TDIQNQQAG----FQALTSYPDDRAHWL
AA_Campylobacterota_WP_010891944.1       NP---------SIDHSPVLNA------------YKLNGDNN------FFSYKLNNEERLG
AA_Chrysiogenetota_WP_183731730.1        --------NYSQLSDTWQKKT-------------PDQRDRG-------YFSRDNSDMVFA
AA_Acidobacteriota_HCZ33114.1            KQ---------KGTDLLGSQA----------SRMEDDAAQG------LPRRMTSGGQTVI
AA_Deinococcota_WP_184108923.1           GR-------VPSLSEVADPAL----RALI---GPDGQVRPG------TREFTVDRQAYSA
AA_Thermodesulfobacteria_RUM89741.1      GMIIT---AAPSVSETEAAED-SVLIS----RIFQAWEKAG-------------------
AA_Fibrobacterota_WP_022637451.1         TRSEN---GLVSLPKAMQSRH-PLLRQSATALAQKTQDKKN------SLALLIENKEIFV
AM_Desulfobacterota_WP_027185430.1       MR--------ESIFSLAETHSSKVLRDIG---KKMVQGETG----FVRLPEFVMGEPAWL
AM_Aproteobac_McpX_WP_014528895.1        MKEYDG---------------------------------EG-A-----------------
AM_Gproteobac_PacA_WP_039291840.1        SKAWQ--------------------------GPTDNFTSS----VVQHDDAILGEQALV
AM_Halobacteriota_WP_011034866.1         GK---------KDLYDFGGEEL----------EKASRDIKNG-IGGHLETADPTTGKTVIL
AM_Spirochaetota_WP_015707342.1          GK---------DLEETLVEGQ-SRLQHID---EIKSAINSG-------EMYISTGKNFYT
AM_Verrucomicrobiota_WP_129046516.1      GK---------PMKDTDPWVV-----------PFLGNLKKG-EAFETESFSRTLNDMTYR
AM_Aproteobac_WP_041812266.1             GQ---------PIGKSDPALD-----------AAKPAIRAG---RSFEQMSVADGQPVKQ
AM_Bacteroidota_WP_092437134.1           GK--------NILEQKNIGAE-----------DQLVDIEKG-------NSLTRIDNGILK
AM_Firmicutes_WP_056043800.1             SKPYLPLPKGESLEELKEQAL-----------------------TIMYTSDPMLGGTVMR
AM_Firmicutes_WP_052635864.1             YN------QDENLKE------------------EILHFKQG--------SHVEYSNGIFD
AM_Gproteobac_PctD_WP_038403940.1        GE----------------------------------------------------------
PU_Gproteobac_R3_WP_124259872.1          DG-------QHFLRDTSGFNQ-----------EMVTKLLMG--QRFSSVSYSATDGERIA
PU_Gproteobac_McpH_WP_047603170.1        GK--------RQLAEQLGADA-------------AKGVMTG-GESLRSSRFSRDGERYLA
PU_Gproteobac_R1_WP_219614703.1          DR--------DNLTQLASGTT----------ANLLTKQAFA------ATQAEVDGQAVIL
PU_Firmicutes_A_R5_WP_131005693.1        EN--------INFFENSSSDL-----------SNSKQLILNNKKEQKSFWYSSEKSKSYI
PU_Actinobacteriota_R8_WP_227113344.1    GH---------ALFDTEEAAV-------------LSEQTRSDRTDVQDFWYRANGENGFL
PU_Cyanobacteria_R9_WP_198537540.1       PP---------QLQRAIAQQR---------LRDSLREQPRG------AFQFRQGGELIYV
PU_Gproteobac_R7_WP_038903150.1          AS---------NIHQQPGLQS------------LATQVLTS---PGGSYRYSLNGENIFL
PU_Gproteobac_R6_WP_185834821.1          ------HPGFDQIQQREGLKT------------LATQILTS---PSVGASYYADGQKVYL
PU_Verrucomicrobiota_R10_WP_110129388.1  RG-------YDSLQEIEGLGE------------RVADLLAG---KTDSLTYKRLGDRRIM
PU_Verrucomicrob_R11_WP_162024566.1      MN-------YQSLQNIPGLND------------VSTDLLAQ---ERTTLTYEREGTTYML
PU_Verrucomicrobiota_R12_WP_220621668.1  TT-------YGNLDAVPGLKK------------DAQSLTSG--TEEQKIRYERDGKTYFL
PU_Campylobacterota_R2_WP_107944080.1    GK--------VNVKDVLGKEK----------FDVLMNKNGG------VIRHFNGTRNLII
PU_Firmicutes_R13_WP_167859577.1         GT---------KLNTILPGAV---SDELF----------AGFANTQVVTYRTAQTGTIDL
PU_Firmicutes_R15_WP_207952809.1         GA---------MISSFVPEEA---FEQML---GQLNETDSGLAKPVVLDYEDQEGKQMDL
PU_Firmicutes_R14_WP_021170906.1         GR---------LAGDFLPAEV---MQQIL---GDM---DNATARPKVLEYQDSQGRIMDL
PU_Gproteobac_R4_WP_199478134.1          KQ--------TDIASLLQDKN-----------------------QVQIVEKSISGKDYYL
```

```
AA_Proteobacteria_NP_252999.1           AFTPI-KGLPSVTWYLALSIDKDKAYAMLSKFRVSA------------------
AA_Cyanobacteria_WP_162544314.1         GFQPI-AGLPGVDWYVAMSIDKGVAFAELSDFRRSTL-----------------
AA_Actinobacteriota_WP_187050447.1      GY----ATNPTTGWRIAGVMEENEVKVRASPM----------------------
AA_Bacteroidota_WP_196119247.1          SY----LTVPNTQWKLVGVNPIN-IDKEVSPIKNRF------------------
AA_Thermotogota_WP_114702178.1          GFSKM-----DNGWIFVTVALSKEVNKRATE-----------------------
AA_Methanobacteriota_WP_056934003.1     GY----KRLSTTGWIVVATVP---------------------------------
AA_Firmicutes_NP_389278.2               VY----QTIGETGWKVGTQFDTDQLMWISDKMNRA-------------------
AA_Desulfobacterota_WP_144684422.1      EV----HTLPDLGWKLVGFIERDEVMEGFRAMMRL-------------------
AA_Chloroflexota_WP_116224856.1         IY----YTSPQLGWKIAAIIPI--------------------------------
AA_Halobacterota_WP_004039109.1         IY----ITSPELGWKIGTFIPVSTIEERINESIRQTL-----------------
AA_Thermoplasmatota_OPX61375.1          VY----YTSTDMGWKIGTFIPNSYLASEIQES----------------------
AA_Spirochaetota_WP_018527208.1         RT----VTLENFGWHLAMAVP---------------------------------
AA_Singergistota_WP_012869308.1         FFSPT-----RSGFVLGVVFPASDLNAMVRSLAIRQ------------------
AA_Verrucomicrobiota_WP_176014353.1     GY----APVASTDWSLAVVLPEADILKQSHCIEETIATESNRLEEEFQLSIHHQ
AA_Planctomycetota_MBG80894.1           LY----EPVQEANWSMGTAIAEDDILAPVHQQ----------------------
AA_Campylobacterota_WP_010891944.1      ACTKV------FAYTACITESADIINKPIYKA----------------------
AA_Chrysiogenetota_WP_183731730.1       SS----TRIPSTQWQLVLFTPRDSFYAPIDPLRN--------------------
AA_Acidobacteriota_HCZ33114.1           GLSRTFSGLPGLHWKLLLSVPE--------------------------------
AA_Deinococcota_WP_184108923.1          VVQPI-ALAPGVQWRVGVYAPVDDFMLGLQRT----------------------
AA_Thermodesulfobacteria_RUM89741.1     ---------MPLDSFVKVSMDSSKWLASLRPLEQGDEA--------FYVG----
AA_Fibrobacterota_WP_022637451.1        NSTP--FHKDNLHWNIVVAMPEAALMGDVRQRQRNSR-----------------
AM_Desulfobacterota_WP_027185430.1      SY----APVSNSDWSMGLVIPEAEMFQGLEGLSRE-------------------
AM_Aproteobac_McpX_WP_014528895.1       ------------------------------------------------------
AM_Gproteobac_PacA_WP_039291840.1       TWQPVTIGNSTEKWYLGI------------------------------------
AM_Halobacteriota_WP_011034866.1        FYEPV----ETGDFAFVLVVPKEEMLAGVADLRER-------------------
AM_Spirochaetota_WP_015707342.1         VYMPIQFSSVTNPWSVAVSIPMAKILANADSIRNY-------------------
AM_Verrucomicrobiota_WP_129046516.1     FGVPVRIGSSSTPWCVSITIRESEVLAGAWKLRNT-------------------
AM_Aproteobac_WP_041812266.1            LFLPVTVAGTETPWSLLVNLPLDKINAPVRELRNAT------------------
AM_Bacteroidota_WP_092437134.1          AFVPVIVGRCPTAWQVSISVPVDYITQEARAQMIYQ------------------
AM_Firmicutes_WP_056043800.1            LLNP--IHIKDQTWYFETIIPKGNMLKDYYKGLSNT------------------
AM_Firmicutes_WP_052635864.1            VYELINIRDIDDKWGIKLSVDQKTIMGSASRLLANQ------------------
AM_Gproteobac_PctD_WP_038403940.1       --------------------PAGSVL----------D-----------------
PU_Gproteobac_R3_WP_124259872.1         AA----SYVPELNLYIVAEVPQAEILGKITQT----------------------
PU_Gproteobac_McpH_WP_047603170.1       LG----LPLRDLNWTLVAEVPESEIYAQMHQA----------------------
PU_Gproteobac_R1_WP_219614703.1         AT----SYIPMLDWYLVAQVPEAEIYAELDKARLH-------------------
PU_Firmicutes_A_R5_WP_131005693.1       VC----QYIPSLKWHLVLENDFTLMIKQLHLQ----------------------
PU_Actinobacteriota_R8_WP_227113344.1   VS----RYLPNLDWFLVVDHDTSQLDAQMARQ----------------------
PU_Cyanobacteria_R9_WP_198537540.1      RT----KRIPELNWTLVVSQPLRVPSGPLWS-----------------------
PU_Gproteobac_R7_WP_038903150.1         NT----RVIPEFGWKLMVEQNSG-------------------------------
PU_Gproteobac_R6_WP_185834821.1         NS----RWVDEFQWYIIVEQKDE-----------------------F-------
PU_Verrucomicrobiota_R10_WP_110129388.1 NC----RYVPELDWYLIVEQSEATMMAPLRQE----------------------
PU_Verrucomicrob_R11_WP_162024566.1     NC----RFIPELNWFLIIEQSEQELLAPIKEQ----------------------
PU_Verrucomicrobiota_R12_WP_220621668.1 NS----RWIPELNWYLMVEQSEDELLSPLRHTLL--------------------
PU_Campylobacterota_R2_WP_107944080.1   GS----KYIPSLDWYLFGEMDEDVLLKDLHT-----------------------
PU_Firmicutes_R13_WP_167859577.1        IS----RPIESADLDIVFAIPRSESVPFLHKIRTNT------------------
PU_Firmicutes_R15_WP_207952809.1        IS----YPLKSTDWKLLFQMRRSESVAFLDTIKLN-------------------
PU_Firmicutes_R14_WP_021170906.1        AY----QSTATTDWKLVFQIPRSESIAILSSVK---------------------
PU_Gproteobac_R4_WP_199478134.1         GA----LWVPMLDRFIVIEVPSEQILSPIYQQ----------------------
```

**Figure S8. Inosine and theophylline do not alter colony morphology and c-di-GMP levels of *P. putida* containing the empty plasmid pBBR1MCS-2_START. A.** Colony morphology and c-di-GMP levels (using c-di-GMP biosensor plasmid pCdrA::*gfp*$^{C}$) in the absence and presence of increasing inosine and theophylline concentrations. The brightness of the colony morphology images has been adjusted to optimize visibility. **B.** c-di-GMP synthesis during growth in liquid cultures of *P. putida* in the absence and presence of different theophylline concentrations. Experiments were conducted under identical condition as those shown in Fig. 7, except that the strain harbored the empty plasmid pBBR1MCS-2_START instead of the R6 expression plasmid pBBR1MCS-2_START_R6.

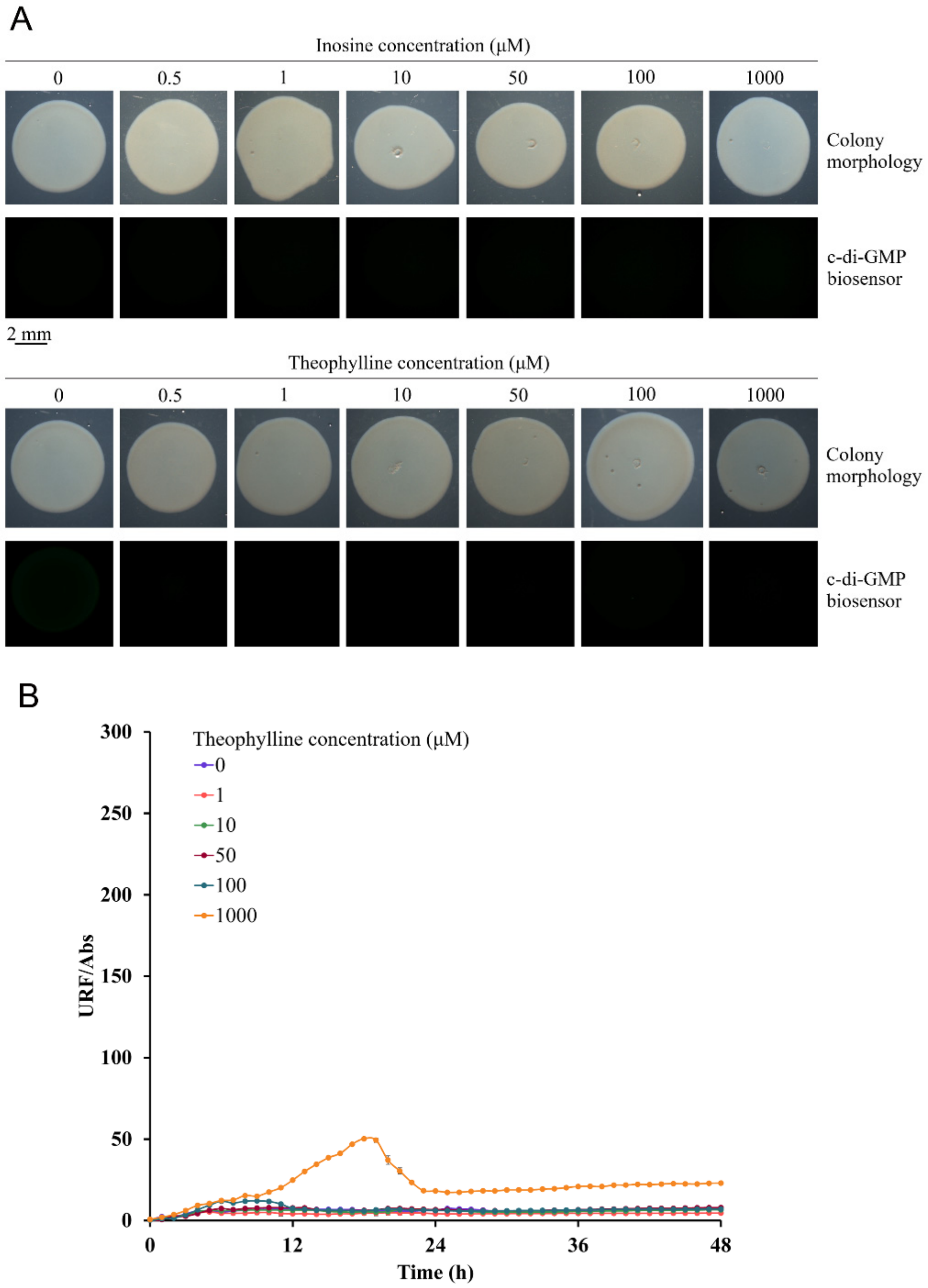

**Table S1. Structural alignment of McpH-LBD with all structures deposited in the protein data bank using the DALI** (3) **algorithm.**

| PDB ID. | Acronym | Z-score | Rmsd[a] (Å) | Identity (%) | Species | Receptor family[b] | Bound ligand | Reference |
|---|---|---|---|---|---|---|---|---|
| 3LIB | mmHK1S-Z3 | 19.1 | 2.9 | 15 | *Methanosarcina mazei* | HK | - | (4) |
| 7PRQ | PctD | 18.8 | 3.0 | 16 | *Pseudomonas aeruginosa* | CR | Choline | (5) |
| 6MNI | PscC | 18.6 | 3.1 | 16 | *Pseudomonas syringae* | CR | Proline | (6) |
| 5LT9 | PctB | 18.4 | 2.8 | 20 | *P. aeruginosa* | CR | L-Arg | (7) |
| 6FU4 | TlpQ | 18.3 | 3.2 | 17 | *P. aeruginosa* | CR | Histamine | (8) |
| 3LIF | rpHK1S-Z16 | 17.9 | 2.7 | 15 | *Rhodopseudomonas palustris* | CR | - | (4) |
| 6D8V | McpX | 17.8 | 2.9 | 22 | *Sinorhizobium meliloti* | CR | 1,1-dimethyl-prolinium | (9) |
| 6PZJ | - | 17.7 | 3.0 | 14 | *Leptospira interrogans* | CR | - | Unpublished |
| 5ERE | - | 17.2 | 2.9 | 11 | *Desulfohalobium retbaense* | Stand-alone sensor domain | - | Unpublished |
| 2ZBB | DctB | 17.2 | 3.0 | 12 | *Escherichia coli* | HK | Malonic acid | Unpublished |
| 3BY9 | DctB | 17.2 | 3.3 | 13 | *Vibrio cholerae* | HK | Succinic acid | (10) |
| 3LIC | soHK1S-Z6 | 16.8 | 3.3 | 16 | *Shewanella oneidensis* | HK | - | (4) |
| 4WY9 | Tlp1 | 16.1 | 3.1 | 12 | *Campylobacter jejuni* | CR | - | (11) |
| 6E0A | TlpA | 16.0 | 3.1 | 12 | *Helicobacter pylori* | CR | - | (12) |
| 3LID | vpHK1S-Z8 | 15.7 | 3.8 | 12 | *Vibrio parahaemolyticus* | HK | - | (4) |
| 4XMQ | Tlp3 (CcmL) | 15.0 | 3.2 | 13 | *C. jejuni* | CR | - | (13) |

[a]Rmsd, root mean square deviation
[b]CR, chemoreceptor; HK, histidine kinase

**Table S2. References for the lifestyle and pathogenicity of species harboring the 15 receptors studied (Table 2).**

| Name | Pathogenicity/characteristics | Refs. |
|---|---|---|
| R1 | Fish pathogen, mainly salmon, important economic losses | (14) |
| R2 | Human pathogen, causes inflammatory bowel disease | (15) |
| R3 | Human pathogen, causes lung infections, high mortality rate | (16) |
| R4 | Halophyte plant isolate, can promote plant growth | (17) |
| R5 | Human pathogen, at times fatal, disrupts gut microbiota | (18) |
| R6 | Human pathogen, causes life-threatening diarrheal disease | (19) |
| R7 | Plant pathogen, in particular maize and rice | (20) |
| R8 | Human pathogen, causes life-threatening bacteremia | (21) |
| R9 | $N_2$-fixing bacterium, isolated from freshwater | (22) |
| R10 | Isolated from a marine saltern | (23) |
| R11 | Efficient degrader of fucoidan, a major algae product | (24) |
| R12 | Isolated from deep sea cold seep | (25) |
| R13 | $N_2$-fixing isolated from orchid roots | (26) |
| R14 | Anaerobic, endospore forming bacterium, isolated from leafs | (27) |
| R15 | Isolated from agricultural soil | (28) |

**Table S3. Presence of theophylline in different samples.**

| Sample type | Comment/reference |
|---|---|
| **Plants and plant extracts** | |
| Black tea | (29, 30) |
| Coffee | (31, 32) |
| Chocolate | (33) |
| Mate | (34) |
| Other plant products like lemon, pummelo or guaran | (35) |
| Bermuda grass | (36) |
| Citrus flowers (*Citrus limon, Citrus maxima, Citrus paradise*) | (37) |
| Different plants | (38) |
| Throughout the plant kingdom | (39) |
| **Human/animals** | |
| *Haliotis discus hannai* (mollusc) | (40) |
| *Anopheles gambiae* (insect) | (41) |
| Human | Many different tissues, cells and excretions (35) |
| Mouse | ST000696 of https://www.metabolomicsworkbench.org |
| Rat | ST000017 of https://www.metabolomicsworkbench.org |
| Baboon | ST002132 of https://www.metabolomicsworkbench.org |
| Goat | ST001968 of https://www.metabolomicsworkbench.org |
| Squirrel | ST000724 of https://www.metabolomicsworkbench.org |
| Pigs | (42) |
| **Microbiomes** | |
| Nasal microbiome | (43) |
| Cervical microbiota | (44) |
| Gut microbiome of insects | (45) |
| **Microorganisms** | |
| *Escherichia soli* | (46) |
| *Staphylococcus aureus* | (47) |
| *Treponema pectinovorum* | Project ST000692 of https://www.metabolomicsworkbench.org |
| *Trypanosoma brucei* | (48) |

**Table S4. Strains and plasmids used.**

| Strains and plasmids | Genotype or relevant characteristics[a] | Ref. |
|---|---|---|
| **Strains** | | |
| *Escherichia coli* BL21(DE3) | F⁻ *ompT gal dcm lon hsdS*$_B$($r_B^- m_B^-$) λ(DE3 [*lacI lacUV5-T7p07 ind1 sam7 nin5*]) [*malB*$^+$]$_{K-12}$($\lambda^S$) | (49) |
| *Pseudomonas putida* KT2440 | Wild type strain | (50) |
| **Plasmids** | | |
| pET28_LBD_McpH | Km$^R$; pET28b(+) derivative containing the DNA fragment encoding McpH-LBD | (51) |
| pET28_LBD_McpH_Y121A | Km$^R$; pET28b(+) derivative containing the DNA fragment encoding McpH-LBD Y121A | This study[a] |
| pET28_LBD_McpH_R129A | Km$^R$; pET28b(+) derivative containing the DNA fragment encoding McpH-LBD R129A | This study[a] |
| pET28_LBD_McpH_W140A | Km$^R$; pET28b(+) derivative containing the DNA fragment encoding McpH-LBD W140A | This study[a] |
| pET28_LBD_McpH_F167A | Km$^R$; pET28b(+) derivative containing the DNA fragment encoding McpH-LBD F167A | This study[a] |
| pET28_LBD_McpH_D169A | Km$^R$; pET28b(+) derivative containing the DNA fragment encoding McpH-LBD D169A | This study[a] |
| pET28_LBD_McpH_D169N | Km$^R$; pET28b(+) derivative containing the DNA fragment encoding McpH-LBD D169N | This study[a] |
| pET28_WP_219614703-LBD | Km$^R$; pET28b(+) derivative containing a DNA fragment encoding WP_219614703-LBD (R1) | This study[a] |
| pET28_WP_107944080-LBD | Km$^R$; pET28b(+) derivative containing a DNA fragment encoding WP_107944080-LBD (R2) | This study[a] |
| pET28_WP_124259872-LBD | Km$^R$; pET28b(+) derivative containing a DNA fragment encoding WP_124259872-LBD (R3) | This study[a] |
| pET28_WP_199478134-LBD | Km$^R$; pET28b(+) derivative containing a DNA fragment encoding WP_199478134-LBD (R4) | This study[a] |
| pET28_WP_131005693-LBD | Km$^R$; pET28b(+) derivative containing a DNA fragment encoding WP_131005693-LBD (R5) | This study[a] |
| pET28_WP_185834821-LBD | Km$^R$; pET28b(+) derivative containing a DNA fragment encoding WP_185834821-LBD (R6) | This study[a] |
| pET28_WP_038903150-LBD | Km$^R$; pET28b(+) derivative containing a DNA fragment encoding WP_038903150-LBD (R7) | This study[a] |
| pET28_WP_227113344-LBD | Km$^R$; pET28b(+) derivative containing a DNA fragment encoding WP_227113344-LBD (R8) | This study[a] |
| pET28_WP_198537540-LBD | Km$^R$; pET28b(+) derivative containing a DNA fragment encoding WP_198537540-LBD (R9) | This study[a] |
| pET28_WP_110129388-LBD | Km$^R$; pET28b(+) derivative containing a DNA fragment encoding WP_110129388-LBD (R10) | This study[a] |
| pET28_WP_167859577-LBD | Km$^R$; pET28b(+) derivative containing a DNA fragment encoding WP_167859577-LBD (R13) | This study[a] |
| pET28_WP_021170906-LBD | Km$^R$; pET28b(+) derivative containing a DNA fragment encoding WP_021170906-LBD (R14) | This study[a] |
| pET28_WP_162024566-LBD | Km$^R$; pET28b(+) derivative containing a DNA fragment encoding WP_162024566-LBD (R11) | This study[a] |
| pET28_WP_207952809-LBD | Km$^R$; pET28b(+) derivative containing a DNA fragment encoding WP_207952809-LBD (R15) | This study[a] |

| | | |
|---|---|---|
| pET28_WP_220621668-LBD | $Km^R$; pET28b(+) derivative containing a DNA fragment encoding WP_220621668-LBD (R12) | This study[a] |
| pBBR1MCS-2_START | $Km^R$; *oriRK2 mobRK2 lacZ* | (52) |
| pBBR1MCS-2_START_R6 | $Km^R$; pBBR1MCS-2_START derivative containing full length coding sequence for WP_185834821 (R6) cloned into NdeI/BamH sites | This study[a] |
| pCdrA::*gfp*[C] | $Ap^R$, $Gm^R$; FleQ dependent c-di-GMP biosensor | (53) |

Ap, ampicillin; Km, kanamycin; Gm, gentamicin.
[a]Gene synthesis and plasmid construction were done by GenScript Inc.

**Table S5. Buffers used for the analysis of the recombinant proteins used in this study.** Buffers were chosen to optimize protein solubility.

| Protein name | Analysis buffer |
|---|---|
| McpH-LBD | 3 mM Tris, 3 mM PIPES, 3 mM MES, pH 6.0 |
| McpH-LBD Y121A | 3 mM Tris, 3 mM PIPES, 3 mM MES, pH 6.0 |
| McpH-LBD R129A | 3 mM Tris, 3 mM PIPES, 3 mM MES, pH 6.0 |
| McpH-LBD W140A | 3 mM Tris, 3 mM PIPES, 3 mM MES, pH 6.0 |
| McpH-LBD F167A | 3 mM Tris, 3 mM PIPES, 3 mM MES, pH 6.0 |
| McpH-LBD D169A | 3 mM Tris, 3 mM PIPES, 3 mM MES, pH 6.0 |
| McpH-LBD D169N | 3 mM Tris, 3 mM PIPES, 3 mM MES, pH 6.0 |
| WP_219614703-LBD (R1) | 3 mM Tris, 3 mM PIPES, 3 mM MES, 150 mM NaCl, 10 % (v/v) glycerol, pH 7.0 |
| WP_107944080-LBD (R2) | 3 mM Tris, 3 mM PIPES, 3 mM MES, 150 mM NaCl, 10 % (v/v) glycerol, pH 8.0 |
| WP_124259872-LBD (R3) | Insoluble |
| WP_199478134-LBD (R4) | 3 mM Tris, 3 mM PIPES, 3 mM MES, 150 mM NaCl, 10 % (v/v) glycerol, pH 8.0 |
| WP_131005693-LBD (R5) | 3 mM Tris, 3 mM PIPES, 3 mM MES, 150 mM NaCl, 10 % (v/v) glycerol, pH 7.0 |
| WP_185834821-LBD (R6) | 3 mM Tris, 3 mM PIPES, 3 mM MES, 150 mM NaCl, 10 % (v/v) glycerol, pH 8.0 |
| WP_038903150-LBD (R7) | Insoluble |
| WP_227113344-LBD (R8) | 3 mM Tris, 3 mM PIPES, 3 mM MES, 150 mM NaCl, 10 % (v/v) glycerol, 1 mM β-mercaptoethanol, pH 6.0 |
| WP_198537540-LBD (R9) | Insufficient protein expression |
| WP_110129388-LBD (R10) | 3 mM Tris, 3 mM PIPES, 3 mM MES, 150 mM NaCl, 10 % (v/v) glycerol, 1 mM β-mercaptoethanol, pH 7.0 |
| WP_162024566-LBD (R11) | 3 mM Tris, 3 mM PIPES, 3 mM MES, 150 mM NaCl, 10 % (v/v) glycerol, 1 mM β-mercaptoethanol, pH 7.0 |
| WP_220621668-LBD (R12) | 3 mM Tris, 3 mM PIPES, 3 mM MES, 150 mM NaCl, 10 % (v/v) glycerol, pH 7.5 |
| WP_167859577-LBD (R13) | 3 mM Tris, 3 mM PIPES, 3 mM MES, 150 mM NaCl, 10 % (v/v) glycerol, pH 7.0 |
| WP_021170906-LBD (R14) | Unfolded protein |
| WP_207952809-LBD (R15) | 3 mM Tris, 3 mM PIPES, 3 mM MES, 150 mM NaCl, 10 % (v/v) glycerol, pH 7.0 |

**Table S6. Sequences of recombinant proteins used in this study**. The hexa-histidine containing extension and mutations to alanine or asparagine are shown in bold face.

| Protein name | Protein sequence (including his-tag) |
|---|---|
| McpH-LBD | **MGSSHHHHHHSSGLVPRGSHM**NRLTDRYLVDTALPASIEAIRNDIERMLG<br>QPLVAAADIAGNTLLRDWLAAGEDPAQAPQFIEYLTAAKQRNHAFTTLFA<br>STETGHYYNENGLDRTLSRSNPKDKWFYGYIDSGAERFINIDIDGATGEL<br>ALFIDYRVEKEGKLVGVAGMGLRMTELSKLIHDFSFGEHGKVFLVRNDGL<br>IQVHPDAAFSGKRQLAEQLGADAAKGVMTGGESLRSSRFSRDGERYLALG<br>LPLRDLNWTLVAEVPESEIYAQMHQ |
| McpH-LBD Y121A | **MGSSHHHHHHSSGLVPRGSHM**NRLTDRYLVDTALPASIEAIRNDIERMLG<br>QPLVAAADIAGNTLLRDWLAAGEDPAQAPQFIEYLTAAKQRNHAFTTLFA<br>STETGH**A**YNENGLDRTLSRSNPKDKWFYGYIDSGAERFINIDIDGATGEL<br>ALFIDYRVEKEGKLVGVAGMGLRMTELSKLIHDFSFGEHGKVFLVRNDGL<br>IQVHPDAAFSGKRQLAEQLGADAAKGVMTGGESLRSSRFSRDGERYLALG<br>LPLRDLNWTLVAEVPESEIYAQMHQ |
| McpH-LBD R129A | **MGSSHHHHHHSSGLVPRGSHM**NRLTDRYLVDTALPASIEAIRNDIERMLG<br>QPLVAAADIAGNTLLRDWLAAGEDPAQAPQFIEYLTAAKQRNHAFTTLFA<br>STETGHYYNENGLD**A**TLSRSNPKDKWFYGYIDSGAERFINIDIDGATGEL<br>ALFIDYRVEKEGKLVGVAGMGLRMTELSKLIHDFSFGEHGKVFLVRNDGL<br>IQVHPDAAFSGKRQLAEQLGADAAKGVMTGGESLRSSRFSRDGERYLALG<br>LPLRDLNWTLVAEVPESEIYAQMHQ |
| McpH-LBD W140A | **MGSSHHHHHHSSGLVPRGSHM**NRLTDRYLVDTALPASIEAIRNDIERMLG<br>QPLVAAADIAGNTLLRDWLAAGEDPAQAPQFIEYLTAAKQRNHAFTTLFA<br>STETGHYYNENGLDRTLSRSNPKDK**A**FYGYIDSGAERFINIDIDGATGEL<br>ALFIDYRVEKEGKLVGVAGMGLRMTELSKLIHDFSFGEHGKVFLVRNDGL<br>IQVHPDAAFSGKRQLAEQLGADAAKGVMTGGESLRSSRFSRDGERYLALG<br>LPLRDLNWTLVAEVPESEIYAQMHQ |
| McpH-LBD F167A | **MGSSHHHHHHSSGLVPRGSHM**NRLTDRYLVDTALPASIEAIRNDIERMLG<br>QPLVAAADIAGNTLLRDWLAAGEDPAQAPQFIEYLTAAKQRNHAFTTLFA<br>STETGHYYNENGLDRTLSRSNPKDKWFYGYIDSGAERFINIDIDGATGEL<br>AL**A**IDYRVEKEGKLVGVAGMGLRMTELSKLIHDFSFGEHGKVFLVRNDGL<br>IQVHPDAAFSGKRQLAEQLGADAAKGVMTGGESLRSSRFSRDGERYLALG<br>LPLRDLNWTLVAEVPESEIYAQMHQ |
| McpH-LBD D169A | **MGSSHHHHHHSSGLVPRGSHM**NRLTDRYLVDTALPASIEAIRNDIERMLG<br>QPLVAAADIAGNTLLRDWLAAGEDPAQAPQFIEYLTAAKQRNHAFTTLFA<br>STETGHYYNENGLDRTLSRSNPKDKWFYGYIDSGAERFINIDIDGATGEL<br>ALFI**A**YRVEKEGKLVGVAGMGLRMTELSKLIHDFSFGEHGKVFLVRNDGL<br>IQVHPDAAFSGKRQLAEQLGADAAKGVMTGGESLRSSRFSRDGERYLALG<br>LPLRDLNWTLVAEVPESEIYAQMHQ |
| McpH-LBD D169N | **MGSSHHHHHHSSGLVPRGSHM**NRLTDRYLVDTALPASIEAIRNDIERMLG<br>QPLVAAADIAGNTLLRDWLAAGEDPAQAPQFIEYLTAAKQRNHAFTTLFA<br>STETGHYYNENGLDRTLSRSNPKDKWFYGYIDSGAERFINIDIDGATGEL<br>ALFI**N**YRVEKEGKLVGVAGMGLRMTELSKLIHDFSFGEHGKVFLVRNDGL<br>IQVHPDAAFSGKRQLAEQLGADAAKGVMTGGESLRSSRFSRDGERYLALG<br>LPLRDLNWTLVAEVPESEIYAQMHQ |
| WP_219614703-LBD (R1) | **MGSSHHHHHHSSGLVPRGSHM**QRSAQQLIETRMFEQELPNLTQRIGKEIE<br>KDLTSVANAARQLANDRFVLDWVARGMPKEQESILIDQLKDMTAQYGLVT<br>ASFADRQSAAYYNQDGFLRNLTPEQDAWFYGYTKSPQDLMLSIFRETNGE<br>VKLFVNFQQLNGRGLAGLAKSLDSMVSMLANFRIGDSGFVFMTDGSGKVK<br>LHPDAARIDRDNLTQLASGTTANLLTKQAFAATQAEVDGQAVILATSYIP<br>MLDWYLVAQVPEAEIYAELDKARLH |
| WP_107944080-LBD (R2) | **MGSSHHHHHHSSGLVPRGSHM**NLYTEKVVKDELPLAVSNVAGEIGYAIDK<br>IINTSYQMTKNDYLLKWIDEGEPKDGLATLFNYNTDLMKAFNLSTAMFVS<br>DKTLNYYTNDKILKQLSKDNPRDSWYFDVKNGKEVNSLNIQVSEATGSLT |

| | |
|---|---|
| | LYVNSKVEKDGKFYGVSAIGMNLDDIVNLVTSKTMGEGSKFLMVDSSGIV<br>KIEKSDRVGKVNVKDVLGKEKFDVLMNKNGGVIRHFNGTRNLIIGSKYIP<br>SLDWYLFGEMDEDVLLKDLHT |
| WP_124259872-LBD (R3) | **MGSSHHHHHHSSGLVPRGSHM**KERAREQDLPTALGEIRSEVLRQIAAPVA<br>LTRSLATNEYILNWEEQGLPEQGAAAWKTYAQTLKNESHAATIAWVSEKT<br>GKYLDENGFSRTVQRSDATDNWFYDFLSKGKQLEIKLGKDKASSVYNLFI<br>DARFDVNGKIGVAALGLSVNELADFIRKQKIGNSGFVYLVSPDGAFVIHR<br>DAALADGQHFLRDTSGFNQEMVTKLLMGQRFSSVSYSATDGERIAAASYV<br>PELNLYIVAEVPQAEILGKITQT |
| WP_199478134-LBD (R4) | **MGSSHHHHHHSSGLVPRGSHM**MESDLKQLKEELLPNRLHSLSSRISEQIS<br>PLINASKLMTNDRFIADWVKKGADESRLPLVAEELNSIKQLSGSDSTFYV<br>VNMKSGLEFLGYDQKFFRTPLADYPYKEFYPNFLAKNKDYELNLQYADQK<br>LYINYRSREMDPTTGKPLVVAGLAIKVDKLIDMVKQLTIGKSGRAMLVTD<br>QGVIQAKGESPAIDLIKQTDIASLLQDKNQVQIVEKSISGKDYYLGALWV<br>PMLDRFIVIEVPSEQILSPIYQQ |
| WP_131005693-LBD (R5) | **MGSSHHHHHHSSGLVPRGSHM**GYQSNTNIFKNDIEHVSTLAAEGIYYQID<br>KLLSEPINVSLTMANDSLLKNFLDGEKEHLNDEEFIYKLQDYLNVYRNKY<br>SYDSVFLVSTKTNNYYHFNGLDRNLSANNSENQWYYTFLKNDDEYSLNVD<br>NDEASNNSITVFVNCKIKDDNGATMGIIGVGLKVNSLQMLLKGYNDKFDV<br>VARLIDDKGFVQLAVDKTGHENINFFENSSSDLSNSKQLILNNKKEQKSF<br>WYSSEKSKSYIVCQYIPSLKWHLVLENDFTLMIKQLHLQ |
| WP_185834821-LBD (R6) | **MGSSHHHHHHSSGLVPRGSHM**HDTLEEQINKDSLPLTSDNIYSEIQQDLI<br>RPIFISSLMAQDTFVREWTLAGEQDPERIIRYLREIQRQYQTISSFYISD<br>RTGHYYHYTGILKQVSESNPNDAWFYRVKNSAPDKNFEVNIDIDTANSQQ<br>TVVFVNYKVFDFENRFLGVIGVGLSSDAVSALVEKYQKRYNRHIYFINEL<br>GEVTLHGSHHPGFDQIQQREGLKTLATQILTSPSVGASYYADGQKVYLNS<br>RWVDEFQWYIIVEQKDEFNHD |
| WP_038903150-LBD (R7) | **MGSSHHHHHHSSGLVPRGSHM**ARHSLFDEISESSLPLTSDNVYSEIQRDL<br>LNPIFISSLMAHDTFVKDWVLSNETDPQAMTRYLREIDRRFNTVVSFFVS<br>NNTHRYYDPEKISHTLLETSPEDKWFFDIRDEKDGDPYDIEIGVDPENRT<br>RMDIFINYKVFDYSGNFIGVTGVGLPVQRVTQLIETYEQRYNRTIYLIDE<br>DGDVMLHSKAFHRASNIHQQPGLQSLATQVLTSPGGSYRYSLNGENIFLN<br>TRVIPEFGWKLMVEQNSGPHDRQLWLTLLKN |
| WP_227113344-LBD (R8) | **MGSSHHHHHHSSGLVPRGSHM**GYQSSRAAFEKDAERTSLLAAEGAAREID<br>NRFAEPIDVSLAMAHDTLLADLLSAEPTRGDDEAYADAVCAYLESYREAF<br>GFDSVFLVSTESNRYYHFNGVDRTLERDNPENTWYFDFLDRDEAYSLNVD<br>NDEATADEITMFVNARILDEQGSTLAIVGVGFRMDDLKELLAGFEARTDT<br>RVRLLADDGTIRASTDPNENGHALFDTEEAAVLSEQTRSDRTDVQDFWYR<br>ANGENGFLVSRYLPNLDWFLVVDHDTSQLDAQMARQF |
| WP_198537540-LBD (R9) | **MGSSHHHHHHSSGLVPRGSHM**ANALAAARRQVVESTLPLTLDALSADLQQ<br>DFVQPILFASAMAANTLLIDWVEQGEQPEAAVQRYLSRVQAQHGATTVFF<br>VSEASRRYYHPTGILKTVSPGSAQDAWFFRLRASASSYEVNLDRDTADPS<br>RTTVFVNYKLLGDGGRFLGAVGLGRSTSQLTRRIQQAERTNGIQVMFLDG<br>RGRILFSPRRGQAPPQLQRAIAQQRLRDSLREQPRGAFQFRQGGELIYVR<br>TKRIPELNWTLVVSQPLRVPSGPLWS |
| WP_110129388-LBD (R10) | **MGSSHHHHHHSSGLVPRGSHM**VSRNNVRKTLAESTLPLTSDNVYSEIQRD<br>LLRPVFIASLMANDTFLRDWAIAGEKDRDAIVRYLHEIKIKYGTVSSFFV<br>SDKTLKYYYAHGLLKTVSEDEPRDEWYFRVREMDEPYEINVDPDMANQDA<br>LTIFINYRVRDYAGNFIGATGVGLTVTKVNRLISRYEAKYDRQIYFVDAS<br>GNVVLRPSNSTMRGYDSLQEIEGLGERVADLLAGKTDSLTYKRLGDRRIM<br>NCRYVPELDWYLIVEQSEATMMAPLRQE |
| WP_162024566-LBD (R11) | **MGSSHHHHHHSSGLVPRGSHM**RSNMRLSITESTLPLTSDNVYSEIQRDLL<br>RPIFISSLMANDTFLRDWALNGEVDINQITKFLHEIKVEYSTISSFFVSD<br>ISHNYYHAQGLLKEVKENEPRDAWYFKARDMDAPYEINVDLDMANQDQMT<br>IFINYRVFDYNDNFIGMTGVGLTVKNVNNLISHYEAKYQRQIYFLNKDGE |

| | |
|---|---|
| | IVLRPSNSPLMNYQSLQNIPGLNDVSTDLLAQERTTLTYEREGTTYMLNC<br>RFIPELNWFLIIEQSEQELLAPIKEQ |
| WP_220621668-LBD (R12) | **MGSSHHHHHHSSGLVPRGSHM**SLPLTADNIYSVIQRDLLRPIFISSMMAN<br>DAFLRDWTINGEVDVKQMQRYLEEIRREYGTVTSFFISEKSRNYYYWGGV<br>LKQVDENEPRDVWYFRVREMEKPFEINVDIDMANNDALTVFVNYRVYDYE<br>GNYIGAAGTGLTVNRVNALIEEYEGRFNREIFFVDREGNIILGPSKGRLT<br>TYGNLDAVPGLKKDAQSLTSGTEEQKIRYERDGKTYFLNSRWIPELNWYL<br>MV |
| WP_167859577-LBD (R13) | **MGSSHHHHHHSSGLVPRGSHM**TQREAVDKLKTKDLLHLADSIAAKVDGQI<br>RKAKETSLLMAEDPNVINWVAGGERDEALGAVVGRKLTLLTSEHGYDNSF<br>LVSAVSNRYWGEGGTVLRTMSRSDPEDAWFYANVDNRKPIEIVLDYAIAR<br>GDTFVFVNALIGGAERALGETGVGLSLKQSAEQFQQFKYGDKSHLWLVDR<br>EGTIYLSDRYEQAGTKLNTILPGAVSDELFAGFANTQVVTYRTAQTGTID<br>LISRPIESADLDIVFAIPRSESVPFLHKIRTNT |
| WP_021170906-LBD (R14) | **MGSSHHHHHHSSGLVPRGSHM**THNAMVDKLKNRDMLYIVQSMSEKIDGRI<br>ERAQETSLLLADDPTVLAWVESGGRDEAAGEIVKTKITDIGKNYDYVKAF<br>VASTVTNQYWEDNKVVKPLSKTSYTDKWFYKALKSGKKIELNIDYDSIHN<br>ETFIFFNTLVGDVRQPVAVAGVALSLGDIAKEFGSYKFGEHSNLWLVDKQ<br>GKIHLADDLEYNGRLAGDFLPAEVMQQILGDMDNATARPKVLEYQDSQGR<br>IMDLAYQSTATTDWKLVFQIPRSESIAILSSVK |
| WP_207952809-LBD (R15) | **MGSSHHHHHHSSGLVPRGSHM**EKEVVNKLKSKDLVRIAESIASKIDGRLQ<br>RAQESSLTMAMDPELIEWLASGEKDQIAGAHVLQKLDNLAKGFDYSNGFI<br>ASRMTGSYWIEGKQMSKILSQTNPADKWFYDSFASQKRMALNIDYNDSRR<br>DTFVFVNVLVGDVESPVGVAGIGLSLKELANNFTNYKYGANSNVWLVDKS<br>GKIYLSDRVEDIGAMISSFVPEEAFEQMLGQLNETDSGLAKPVVLDYEDQ<br>EGKQMDLISYPLKSTDWKLLFQMRRSESVAFLDTIKLN |

**Table S7. Data collection and refinement statistics**. Statistics for the highest-resolution shell are shown in parentheses.

| Ligand | Uric Acid |
|---|---|
| PDB ID | 8BMV |
| Beam Line | XALOC (ALBA) |
| Space group | P 1 21 1 |
| Unit cell a, b, c (Å) | 32.581 124.322 57.201 |
| Resolution (Å) | 32.15 - 1.95 (2.02 - 1.95) |
| Unique reflections | 31376 (3258) |
| Multiplicity | 2.9 (3.1) |
| Completeness (%) | 96.01 (99.48) |
| $I/\sigma_I$ | 13.07 (1.73) |
| Wilson B-factor | 36.6 |
| $R_{merge}$ (%) | 5.17 (70.7) |
| CC(1/2 | 0.998 (0.678) |
| CC | 1 (0.899) |
| **Refinement** | |
| $R_{work}/R_{free}$ (%) | 18.27 / 23.12 |
| $CC_{work}/CC_{free}$ | 0.97 / 0.91 |
| No. atoms | 4177 |
| Protein | 4037 |
| Ligands | 24 |
| Solvent | 116 |
| B-factor (Å$^2$) | 47.50 |
| R.m.s deviations | |
| Bond lengths (Å) | 0.009 |
| Bond angles (°) | 1.01 |
| Ramachandran (%) | |
| Favoured (%) | 97.55 |
| Outliers (%) | 0.00 |
